# Measuring Cell Type Resolved Aging Through Long-Read Sequencing

**DOI:** 10.64898/2026.11.22.768438

**Authors:** Yinjie Wu, Harlan Stevens, Alec Eames, Mahdi Moqri, Vadim N. Gladyshev

**Affiliations:** Division of Genetics, Department of Medicine, Brigham and Women’s Hospital, Harvard Medical School, Boston, MA, USA; Department of Epidemiology and Biostatistics, Harvard T.H. Chan School of Public Health, Boston, MA, USA; Department of Genetics, Harvard Medical School, Boston, MA, USA

## Abstract

Epigenetic clocks based on bulk blood DNA methylation are powerful tools for quantifying biological aging, but bulk measurements conflate cell-autonomous epigenetic changes with age-related shifts in blood cell composition. Here, we develop LongReadClock, a set of cell-type-resolved epigenetic aging clocks constructed from cohort-scale long-read whole-genome sequencing of bulk blood without physical cell sorting. Individual sequencing reads were classified using seven target-specific blood-cell models using methylation-based log-likelihood-ratio scoring against a purified-cell atlas. LongReadClock achieved mean absolute errors of 2.8–5.7 years on held-out samples across bulk blood and digitally resolved cell types. Covariate-adjusted analyses identified associations that differed across cell types, including lower mean epigenetic age deviation (EAD) in males in B-cell and NK-cells, associations of systolic blood pressure and resting heart rate with myeloid EAD, and higher NK-cell EAD in participants with asthma. A Fisher-information screen was used to prioritize age-informative genomic windows and to explore read-level methylation patterns at the *ELOVL2* locus. These results support cell-type-resolved epigenetic age inference from cohort-scale long-read methylation data from standard bulk blood samples.

## Introduction

Aging is a dominant risk factor for many of the chronic diseases that account for the greatest burden of human morbidity and mortality, including cardiovascular disease, cancer, type 2 diabetes, and neurodegenerative disease [1, 2, 3, 4, 5, 6, 7]. This relationship is reflected in contemporary mortality patterns: in the United States, heart disease and cancer remained the two leading causes of death in 2024, with most other leading causes similarly age-associated [8]. As the global population continues to age, the burden of these conditions is projected to rise substantially, making a deeper mechanistic understanding of the aging process an increasingly urgent scientific and public health priority [9].

Because individuals of the same chronological age can differ substantially in their rate of biological aging, chronological age alone is an imperfect measure of biological aging and disease risk. This has motivated the development of molecular biomarkers that more directly capture biological aging trajectories, among the most widely used of which are DNA methylation-based epigenetic clocks [10]. These clocks exploit reproducible age-associated changes in CpG methylation across the genome to estimate biological age [11, 12]. While early clocks were trained to predict chronological age, subsequent generations were designed to better capture physiological deterioration and health risk, improving their ability to predict mortality, morbidity, and age-related disease outcomes [13, 14]. Epigenetic clocks have since become central tools in human aging research, and epigenetic age deviation (EAD), defined as the extent to which clock-predicted age deviates from chronological age, has emerged as a reproducible correlate of aging-related outcomes across cohorts.

Most established epigenetic clocks were developed from bulk tissue measurements, which in blood introduces an important interpretational challenge. Whole-blood methylation profiles reflect two distinct sources of age-associated signal: an intrinsic component capturing within-cell-type epigenetic changes within individual cell types, and an extrinsic component driven by shifts in blood cell type composition across the lifespan [15, 16]. Because these two processes are conflated in bulk measurements, it is difficult to determine whether an observed clock signal reflects epigenetic aging within cells, compositional remodeling of the hematopoietic system, or some combination of both. Their conflation has measurable consequences. Epigenetic age estimates from the same individual have been shown to vary systematically within a single day, with predicted age differing by several years depending on time of sampling, an effect partially but not fully explained by diurnal fluctuations in blood cell composition [17].

Prior efforts have addressed this limitation through two main strategies. Statistical deconvolution methods adjust bulk methylation profiles for estimated cell type proportions, but these fractions are themselves inferred from the same bulk methylation data, and the adjusted signals remain difficult to interpret at cell-type-specific resolution [18, 19, 20, 21]. More direct sorted-cell approaches have demonstrated that cell-type-specific epigenetic aging signals can be measured within defined immune compartments, but they require specialized cell isolation and methylation profiling workflows that are difficult to scale to large cohorts [22]. These considerations motivate approaches for resolving cell-type-specific methylation aging signals more directly at population scale, complementing bulk-level inference with greater biological resolution.

Recent methodological advances have begun to move beyond conventional bulk-tissue epigenetic clocks toward more resolved views of methylation aging. The scAge framework demonstrated that probabilistic age inference is feasible even in highly sparse single-cell methylation data, providing a conceptual basis for estimating epigenetic age when only a small and variable subset of age-informative CpGs is observed in each profile [23]. More recently, LongReadAge extended this probabilistic strategy to long-read sequencing (lrWGS) and introduced a method for profiling epigenetic aging at cell-type resolution from bulk samples without physical cell sorting [24]. Together, these studies suggest that sparse methylation observations do not need to preclude biologically informative age inference. However, robust approaches for resolving cell-type-specific aging signals from cohort-scale long-read blood methylation data remain limited.

The All of Us Research Program provides a large-scale setting to address this gap [25, 26]. As part of its long-read whole genome sequencing initiative, the program has generated lrWGS data across a large and ancestrally diverse cohort, with genome-wide CpG methylation calls available as a byproduct of basecalling [27, 28]. This dataset enables systematic evaluations of cell-type-resolved epigenetic age inference from long-read blood methylation data at cohort scale. In this project, we develop and evaluate an approach for cell-type-specific epigenetic age inference from these data, combining methylation-based digital sorting of reads into blood cell populations with likelihood-based age prediction using age-associated CpG reference models. We focus first on broad hematopoietic compartments, particularly myeloid and lymphoid lineages, and then extend the analysis to finer immune cell subtypes. Through systematic evaluation of pooled bulk, technology-specific, and cell-type-matched reference schemes, we characterize the conditions under which cell-type resolution yields biologically interpretable aging signals beyond what bulk clocks provide. Finally, we explore the limits of sparse single-read age inference by evaluating individual long reads spanning the highly informative ELOVL2 locus to characterize the information-theoretic constraints that govern the age information available in a single sequencing molecule.

## Results

### Development and application of LongReadClock

Building on the probabilistic age inference strategy of scAge and LongReadAge [23, 24], we developed LongReadClock, a set of cell-type-resolved epigenetic aging clocks for blood constructed from cohort-scale long-read methylation data from the All of Us Research Program. The analytical workflow comprised four main steps (Figure 1). First, CpG methylation calls were extracted from whole-genome long-read sequencing data across all participants, producing genome-wide bulk methylation profiles. Second, differentially methylated marker regions were identified from a purified-cell WGBS atlas, and individual sequencing reads were digitally classified using seven target-specific models for blood cell populations (myeloid, lymphoid, T cell, B cell, NK cell, monocyte, and granulocyte) using a likelihood-ratio scoring approach against reference methylation patterns derived from a purified-cell WGBS atlas [29], without physical cell sorting. Third, CpG-specific linear age associations were estimated within each cell-type compartment from the training participants, constructing separate reference models for bulk blood and each of the seven cell types. Fourth, age was inferred for each participant and cell type by maximizing a Binomial log-likelihood over a discrete age grid, selecting the top *K* = 1,000 age-informative CpGs ranked by their training-set *t*-statistic.

**Figure 1.**
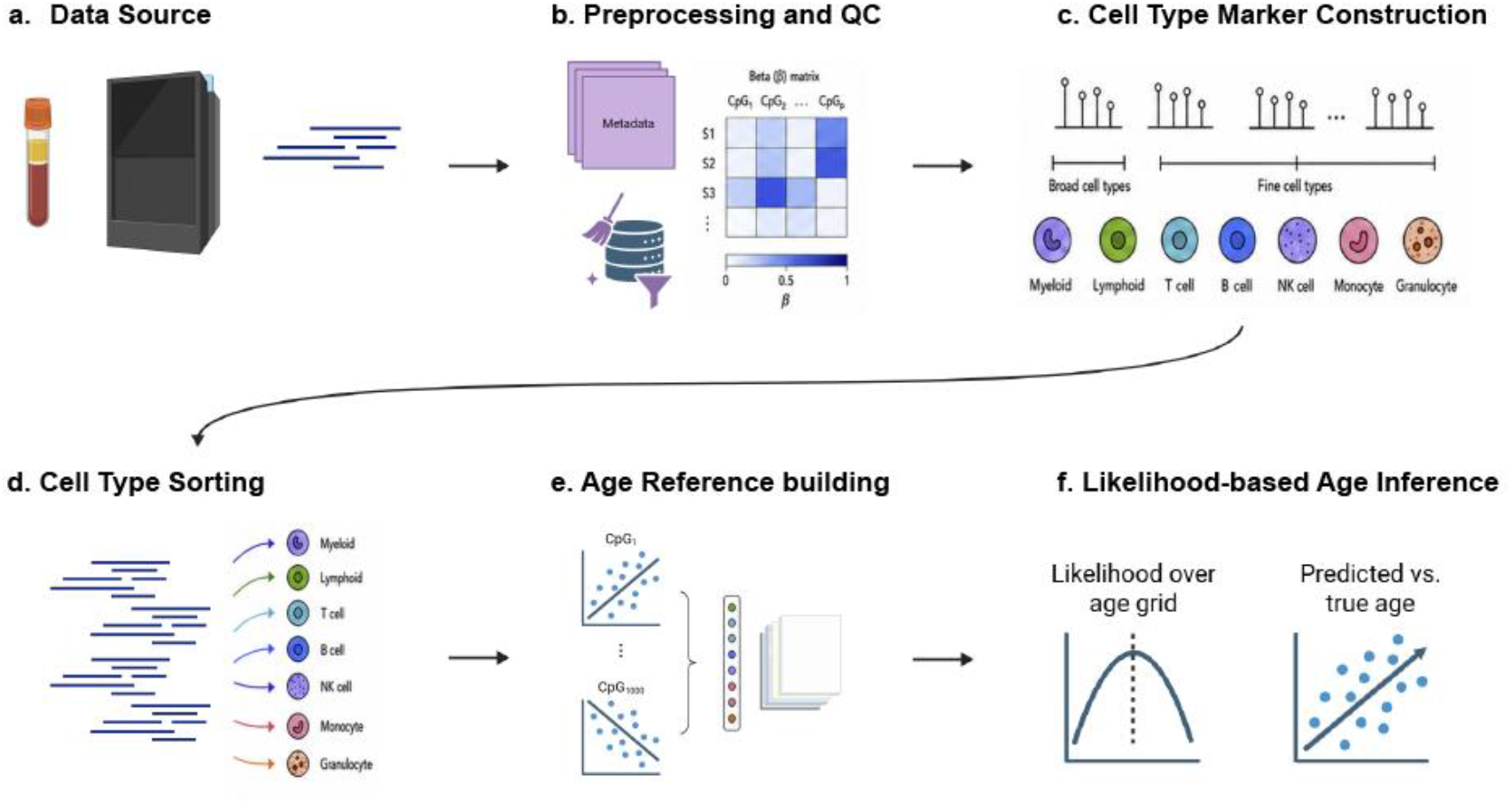
LongReadClock analytical workflow. (a) Cohort-scale long-read blood methylation data and participant metadata were obtained from the All of Us Research Program. (b) Long-read methylation data were processed with wgbstools, linked with blood-draw age and participant metadata, and screened for sample- and batch-level quality before downstream analysis. (c) Differentially methylated marker regions were derived from purified-cell reference methylation profiles for broad myeloid and lymphoid lineages and five finer blood cell types. (d) Individual long-read methylation patterns were digitally assigned to myeloid, lymphoid, T cell, B cell, NK cell, monocyte, and granulocyte outputs by target-versus-background likelihood-ratio scoring. (e) Bulk and digitally sorted training data were used to fit CpG-specific age-reference models and select age-informative autosomal CpGs. (f) Age was inferred from observed informative CpGs by maximising the likelihood across a discrete age grid.

The All of Us long-read resource included 2,842 platform-specific sequencing records from 2,778 unique participants with available age metadata. The difference between these counts reflects participants represented by more than one sequencing record. Data were generated across multiple sequencing centers using Oxford Nanopore Technology (ONT), PacBio Revio, and PacBio Sequel II/IIe instruments (Supplementary Figure S1).

### Platform-specific methylation structure motivates platform selection

We examined platform- and batch-related structure across the full lrWGS dataset to identify subsets suitable for downstream clock construction and evaluation. Principal component analysis of bulk methylation profiles revealed strong separation by sequencing platform and source, indicating substantial technical heterogeneity across batches (Figure 2a). This structure was most pronounced for ONT, Revio, Sequel IIe, and the Controlled Tier v7 samples, suggesting that platform-specific methylation characteristics dominated global methylation variation in the cohort.

**Figure 2.**
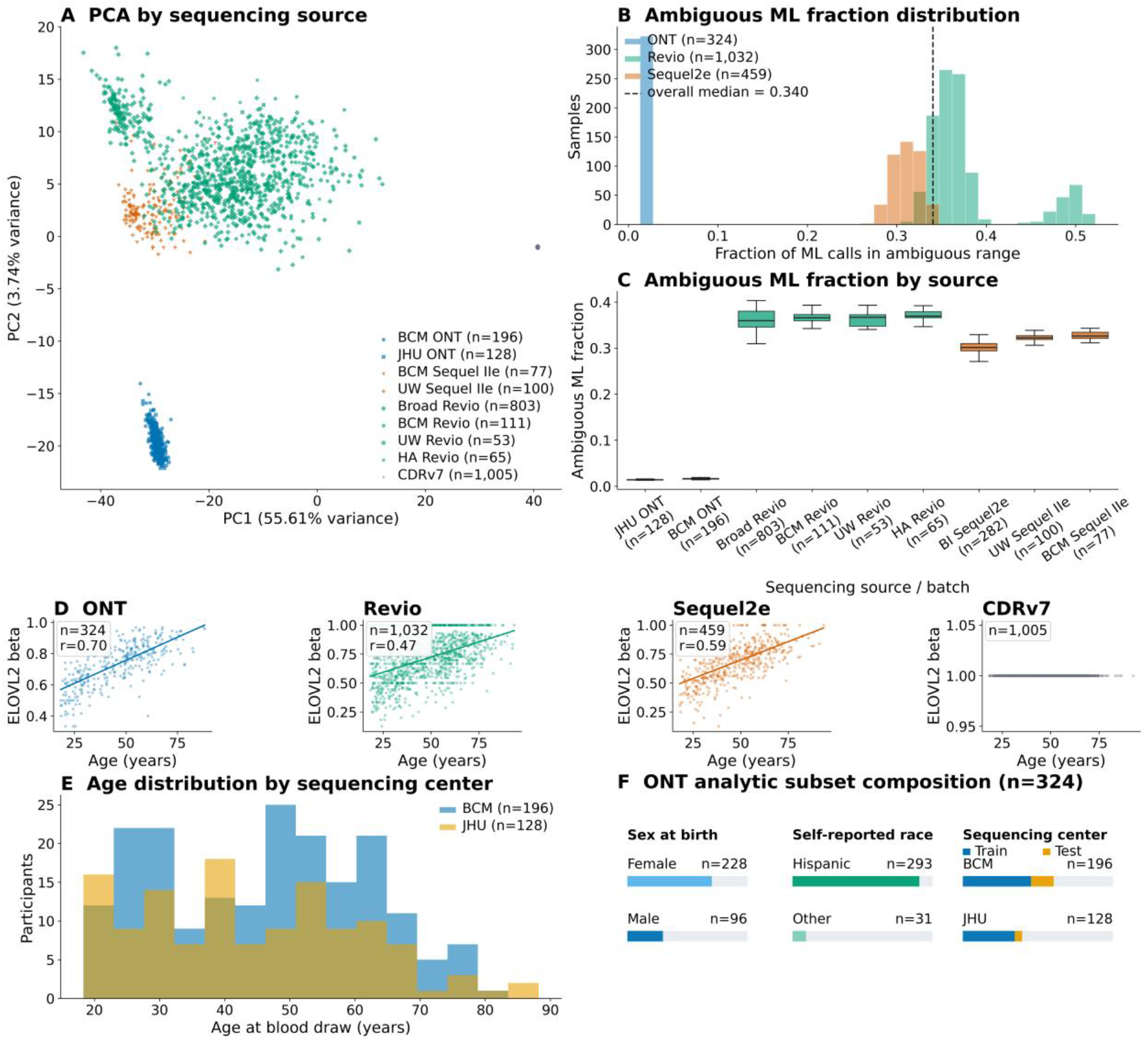
Platform-specific methylation structure and quality assessment across the All of Us long-read methylation cohort. (a) Principal component analysis of bulk methylation profiles, colored by platform and by sequencing source, showing strong batch and platform effects. (b) Distribution of the ambiguous ML fraction across major platforms, where ambiguous ML fraction is defined as the proportion of CpG calls with ML scores between 51 and 204, corresponding to methylation probabilities between 0.20 and 0.80. (c) Ambiguous ML fraction stratified by sequencing source, showing that platform- and batch-related methylation differences are largely driven by methylation calling confidence. (d) Methylation at a canonical age-associated CpG in the ELOVL2 locus (chr6:11044644) plotted against chronological age for each major platform. ONT samples show the strongest age correlation and least apparent noise, whereas Controlled Tier v7 samples show no informative variation. (e) Demographic composition of the ONT analytic subset, including age distribution, age by sequencing center, sex at birth, self-reported race and ethnicity, sample counts by sequencing center, and age distribution by sequencing center.

To investigate the source of this batch effect, we quantified methylation calling uncertainty using the ambiguous ML fraction, defined as the proportion of CpG calls with ML scores between 51 and 204, corresponding to methylation probabilities between 0.20 and 0.80. This analysis showed marked differences across platforms (Figure 2b–c). ONT samples had consistently low ambiguous ML fractions, indicating high-confidence bimodal methylation calls, whereas Revio and Sequel IIe samples showed substantially higher ambiguity. The Controlled Tier v7 samples were particularly problematic, with 1,005 samples showing essentially no usable methylation signal, consistent with their complete loss of informative variation in downstream analyses.

We further evaluated methylation quality at a well-established age-associated CpG in the ELOVL2 locus (chr6:11,044,644; region corresponding to cg16867657). Among the major platforms, ONT showed the clearest age-associated trend and the least apparent noise, while Revio and Sequel IIe displayed weaker and more dispersed relationships; the Controlled Tier v7 samples again showed no informative variation (Figure 2d). Together, these results indicate that the major batch effects observed in the cohort were driven largely by differences in methylation calling confidence and data quality rather than biological variation. Because ONT samples showed the highest-confidence methylation calls and the most reliable age-associated signal, we selected the 324 ONT sequenced participants from Baylor College of Medicine and Johns Hopkins University as the primary analytic cohort for downstream clock construction and phenotype association analyses. These participants spanned a broad adult age range and were of self-reported Hispanic/Latino ethnicity (Figure 2e–f).

### Digital cell-type sorting enables accurate cell-type-specific age prediction

To assess whether digital sorting could recover biologically meaningful cell-type-specific methylation structure, we compared averaged ONT cell-type-predicted methylation profiles against purified-cell reference profiles from the Human Cell Type DNA Methylation Atlas [29] at lineage-discriminating marker CpGs. Representative profiles demonstrated that reads assigned to each cell type recapitulated target-specific methylation patterns (Figure 3j). Quantitative comparisons showed lower mean absolute methylation differences from matched than background purified-cell reference profiles for the myeloid and lymphoid lineages (Figure 3k), a pattern that extended across all seven target models (Supplementary Figure S2).

**Figure 3.**
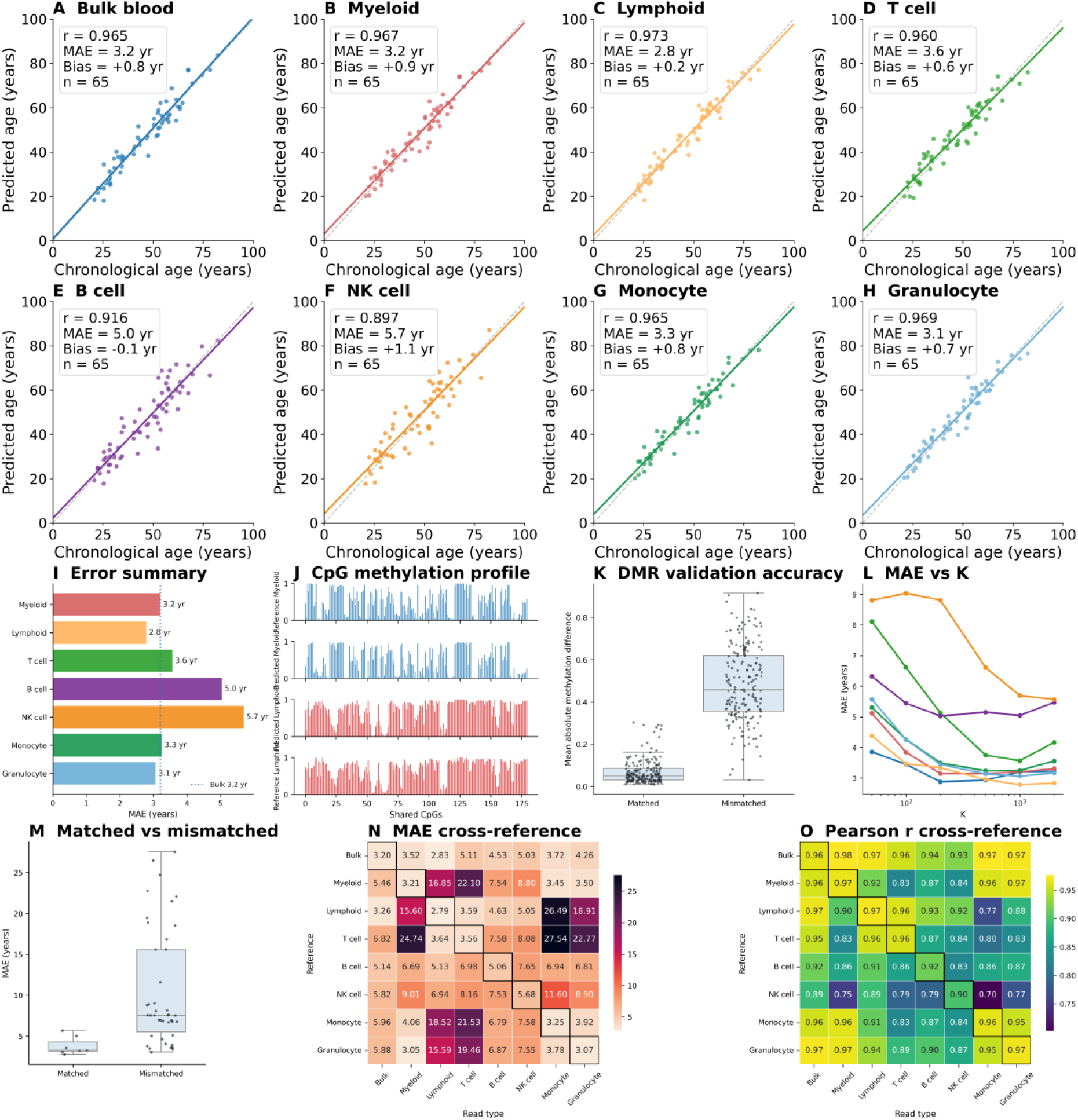
Cell-type-specific age prediction and validation of digital sorting in the ONT subset. (a–h) Predicted versus chronological age on the held-out ONT test set for bulk blood and cell-type-specific self-reference clocks, using K = 1000 age-informative CpG sites. Panels show bulk blood, myeloid, lymphoid, T cell, B cell, NK cell, monocyte, and granulocyte predictions. (i) Summary of mean absolute error (MAE) across bulk and cell-type-specific clocks. (j) Example methylation profiles from the purified-cell methylation atlas and averaged long-read methylation profiles for reads predicted to originate from myeloid and lymphoid compartments, shown for a random set of CpGs overlapping lineage-discriminating marker sites. (k) Mean absolute methylation difference between predicted myeloid and lymphoid profiles and the corresponding myeloid and lymphoid references, showing that predicted profiles more closely match their lineage-consistent reference. (l) Clock performance as a function of the number of age-informative CpG sites (K) for ONT self-reference models. (m–o) Cross-reference evaluation of cell-type-specific clocks, showing that matched reference and read-type pairs yield lower MAE and higher age-prediction correlation than mismatched lineage comparisons.

Having established the biological validity of the sorting procedure, we then evaluated LongReadClock prediction accuracy using ONT self-reference models on the held-out test set (*n* = 65). With *K* = 1,000 age-informative CpG sites, bulk blood age prediction showed strong concordance with chronological age (MAE = 3.2 yr, *r* = 0.965; Figure 3a). Cell-type-specific clocks achieved comparable or better performance across most compartments (Figure 3b–h): the lymphoid clock achieved the lowest MAE among all clocks (2.8 yr, *r* = 0.973), while myeloid (3.2 yr), T cell (3.6 yr), monocyte (3.3 yr), and granulocyte (3.1 yr) clocks all performed within the bulk baseline range. B cell and NK cell clocks showed modestly reduced accuracy (MAE = 5.0 yr and 5.7 yr, respectively), consistent with the sparser methylation coverage expected for these lower-abundance populations in digitally sorted long-read data, though both retained meaningful age correlation (*r* = 0.916 and *r* = 0.898). Taken together, the error summary across all compartments confirmed that cell-type-specific clocks are broadly competitive with bulk blood prediction, with most cell types falling at or below the bulk MAE of 3.2 yr (Figure 3i).

We examined how prediction accuracy scaled with the number of age-informative CpG sites *K*. Performance improved substantially at small panel sizes before plateauing above approximately *K* = 500–1,000 across all clocks (Figure 3l), supporting the choice of *K* = 1,000 for all subsequent analyses. Cross-reference evaluation further confirmed the specificity of cell-type-resolved aging signals. When clocks were applied using a mismatched reference lineage, prediction error increased substantially relative to the matched condition (Figure 3m). This pattern was consistent across all pairwise combinations of reference and read types: the MAE heatmap showed pronounced off-diagonal elevation, with the largest errors arising when lymphoid reads were evaluated against myeloid references and vice versa (Figure 3n). Correspondingly, Pearson correlation was highest along the diagonal and degraded systematically for mismatched pairs (Figure 3o), confirming that cell-type-specific methylation aging signals are sufficiently distinct to support lineage-resolved clock construction and that cross-lineage application meaningfully reduces predictive accuracy.

We benchmarked LongReadClock against elastic-net regression on ONT bulk methylation profiles. LongReadClock achieved comparable age-prediction accuracy while retaining the key advantage of a likelihood framework that accommodates sparse read-level and cell-type-resolved data by evaluating only observed informative CpGs. Five-fold cross-validation showed no evidence of substantial overfitting across bulk and sorted cell-types (Supplementary Figure S3). Residual analyses revealed modest regression-to-the-mean effects at the extremes of the age range, consistent with the bounded nature of the likelihood grid estimator (Supplementary Figure S4).

We next evaluated whether ONT-trained LongReadClock references could generalize to independent long-read platforms. PacBio methylation profiles contained a higher fraction of intermediate-confidence methylation calls, and retaining these ambiguous calls substantially degraded age prediction. We therefore excluded calls with methylation probability values between 0.20 and 0.80 before evaluating. This filtering improved call reliability but also reduced effective CpG coverage, especially in digitally sorted low-abundance cell types.

Using this filtered call set, we compared references trained on ONT, PacBio Sequel IIe, PacBio Revio, or all long-read samples across ONT, Sequel IIe, Revio, and combined test sets. The ONT-trained reference generalized well to PacBio samples and remained competitive even when evaluated on PacBio test sets, indicating that the age-associated methylation signal learned from ONT data transferred across long-read platforms (Supplementary Figure S5). Lower performance in some PacBio comparisons was most apparent in low-abundance sorted compartments, consistent with reduced effective coverage and lower methylation-call confidence rather than failure of the ONT-derived age signal itself. Together with the ONT cross-validation results, these findings support the ONT-trained references as the primary LongReadClock models for downstream epigenetic age deviation analyses.

### Association of age-deviation residuals with phenotypes

To evaluate whether LongReadClock captures biologically meaningful variation in epigenetic aging, we tested associations between out-of-fold EAD residuals and 31 phenotypes spanning demographic, lifestyle, clinical, laboratory, medication, and physiological domains in the ONT cohort (*n* = 324). All models included quadratic age adjustment and, where appropriate, an adaptive sex covariate (Supplementary Figure S6; Methods), and were restricted to autosomal CpG sites (chr1–22). Full association results are provided in Supplementary Table S1, with phenotype coverage summarized in Supplementary Table S2.

Age-adjusted EAD residuals across the eight clocks were moderately to strongly correlated (Pearson *r* = 0.35–0.92; Figure 4a), yet each retained substantial clock-specific variation. Between 29% and 88% of cell-type EAD variance was not explained by bulk EAD with the B cell (88%) and NK cell (83%) clocks showing the greatest divergence from bulk (Figure 4b). Hierarchical clustering separated the myeloid-lineage clocks (myeloid, monocyte, and granulocyte) from the lymphoid-lineage clocks (lymphoid, T cell, B cell, and NK cell), consistent with lineage-resolved biological signal.

**Figure 4.**
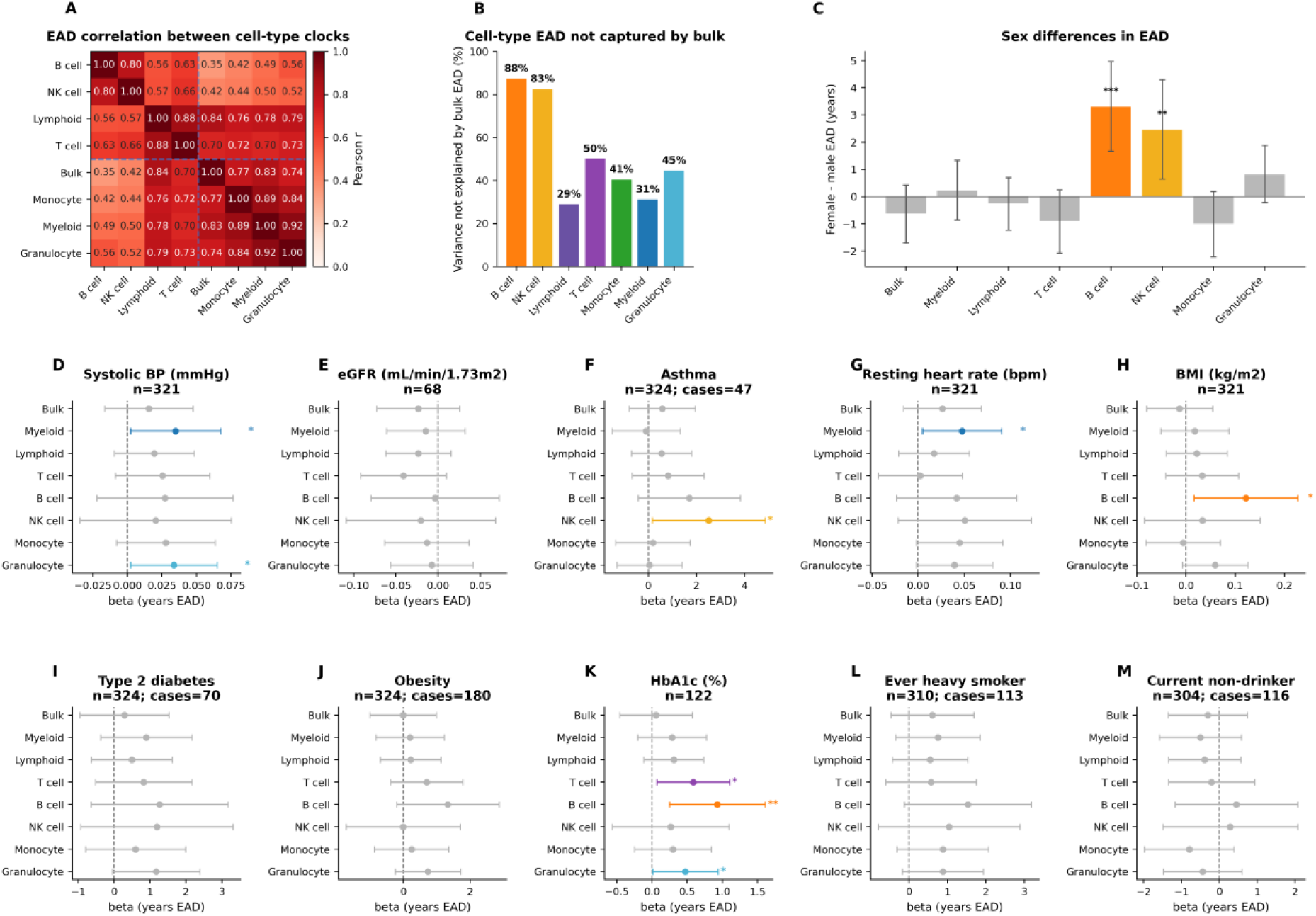
Association of epigenetic age deviation (EAD) residuals with phenotypes across bulk blood and seven cell-type-specific clocks. (a) EAD residual correlation across clocks. (b) Percentage of cell-type EAD variance not explained by bulk EAD. (c) Sex differences in EAD, shown as male–female difference with 95% confidence intervals. (d–m) Forest plots of association coefficients across all eight clocks for systolic blood pressure (mmHg), eGFR (mL/min/1.73 m^2^), asthma, resting heart rate (bpm), BMI (kg/m^2^), type 2 diabetes, obesity, HbA1c (%), ever heavy smoking, and current non-drinking. For continuous phenotypes, n indicates the complete-case sample size. For binary phenotypes, panel titles show total complete-case n and number of cases. Asterisks indicate nominal significance: ^∗^p < 0.05, ^∗∗^p < 0.01, ^∗∗∗^ p < 0.001.

Next, we examined associations between EAD and demographic and clinical phenotypes across the eight clocks. According to these data, after adjustment for linear and quadratic chronological age, sex-associated EAD estimates differed across immune compartments (Figure 4c). Lower mean EAD was estimated in males relative to females for the B cell clock (*β* = −3.32 yr, 95% CI [−4.97, −1.68], *p* = 0.0001) and the NK cell clock (*β* = −2.48 yr, *p* = 0.008). Estimates for most remaining clocks instead pointed toward lower mean EAD in females, but did not reach statistical significance (Supplementary Figure S6).

Among physiological phenotypes, higher systolic blood pressure was associated with higher myeloid (*β* = +0.035 yr/mmHg, *p* = 0.036) and granulocyte (*β* = +0.034 yr/mmHg, *p* = 0.036) EAD, and higher resting heart rate was associated with higher myeloid EAD (*β* = +0.048 yr/bpm, *p* = 0.030). These associations did not reach statistical significance in bulk blood or the lymphoid clocks (Figures 4d,g). BMI was positively associated with B cell EAD (*β* = +0.12 yr per kg/m^2^, *p* = 0.025; Figure 4h), with no statistically significant association in the remaining clocks. Asthma was associated with higher NK cell EAD (*β* = +2.51 yr, 95% CI [+0.17, +4.86], *p* = 0.037; Figure 4f).

eGFR showed negative estimated associations with EAD across all eight clocks, although none reached statistical significance (all *p* ≥ 0.118; Figure 4e). The largest estimate was observed for the T cell clock (*β* = −0.041 yr per mL/min/1.73 m^2^, 95% CI [−0.092, 0.010], *p* = 0.118). HbA1c showed positive associations with T cell (*β* = +0.58 yr per percentage point, *p* = 0.028), B cell (*β* = +0.93 yr per percentage point, *p* = 0.009), and granulocyte EAD (*β* = +0.47 yr per percentage point, *p* = 0.049; Figure 4k).

Several additional candidate phenotypes did not reach statistical significance. Type 2 diabetes showed a positive estimate for granulocyte EAD (*β* = +1.17 yr, *p* = 0.061). Obesity, defined as a documented obesity diagnosis or BMI of at least 30 kg/m^2^, showed a positive B cell estimate (*β* = +1.33 yr, *p* = 0.090), but did not reach statistical significance. Ever heavy smoking was associated with positive estimates across clocks, although none reached statistical significance. Full association estimates, complete-case sample sizes, confidence intervals, nominal *p* values, and Benjamini–Hochberg adjusted *q* values are provided in Supplementary Table S1. Phenotype coverage is summarized in Supplementary Table S2.

### Analyses of single-molecule age inference

To complement whole-genome bulk age estimation, we investigated whether chronological age could be inferred from individual long reads restricted to a single targeted genomic window. This setting is relevant for applications where only a subset of the genome is accessible, such as targeted sequencing or spatially resolved epigenomics.

Ranking all 286,351 genomic windows (10 kb windows, 2.5 kb step) by summed Fisher information for age estimation revealed a characteristic distribution in which informativeness was highest in windows containing 10–30 eligible CpGs and declined with increasing CpG density (Figure 5a). Windows densely packed with CpGs were predominantly composed of weaker age predictors, while the small subset of windows containing highly informative sites stood out clearly. The *ELOVL2* window (chr6:11,039,644–11,049,644; 23 eligible CpGs; starred in Figure 5a) ranked among the top windows genome-wide with a summed Fisher information of 0.019, reflecting the exceptional per-site informativeness of the CpG cluster at this locus and motivating its selection as the primary region for single-molecule evaluation.

**Figure 5.**
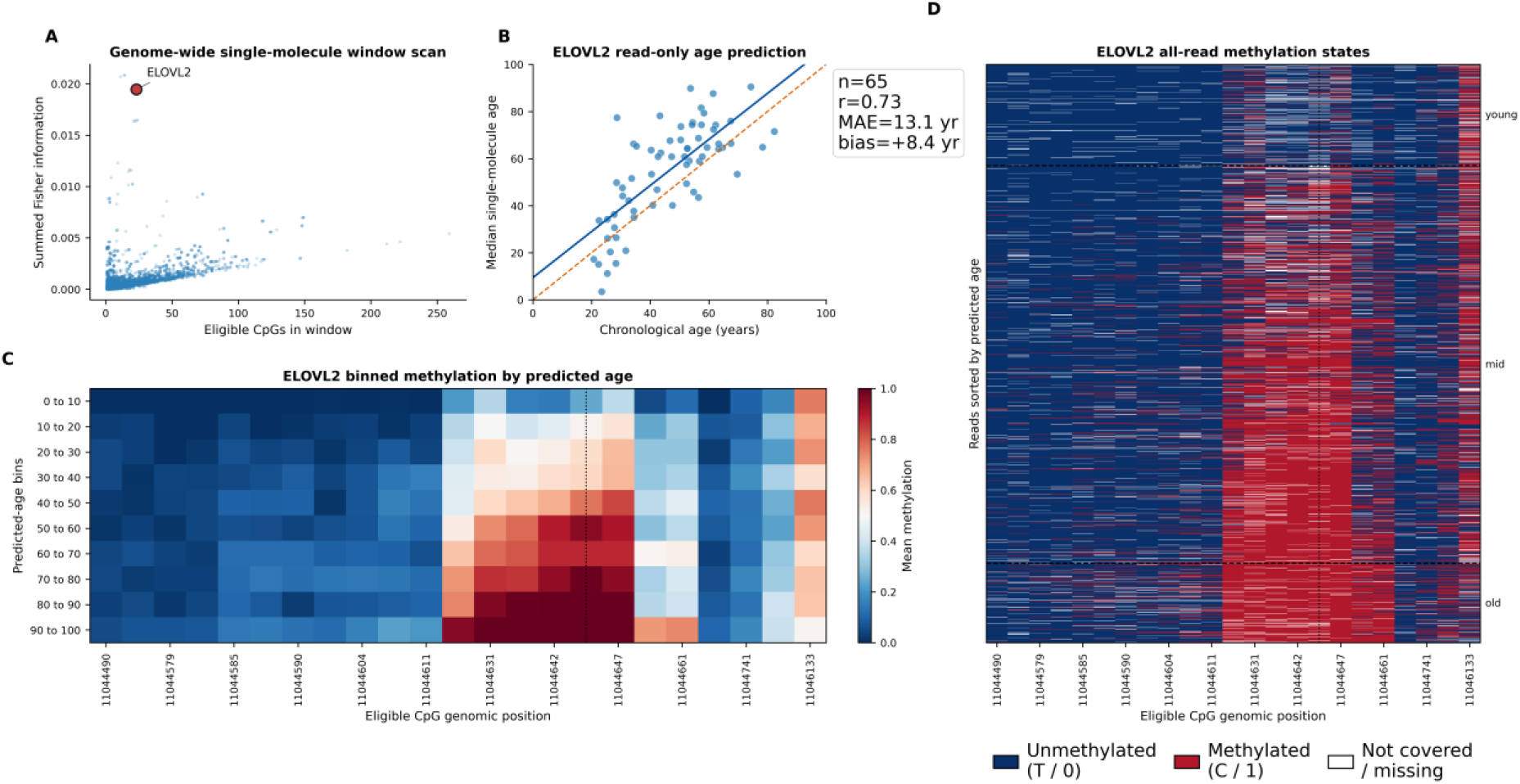
Single-molecule age inference at the ELOVL2 locus. (a) Summed Fisher information versus number of eligible CpGs for all 286,351 genomic windows (10 kb, 2.5 kb step). Each point is one window; the starred point marks the ELOVL2 window. (b) Sample-level weighted-median single-read predicted age from the ELOVL2 window versus chronological age for 20 held-out ONT samples. Blue line = ordinary least squares fit; dashed orange = identity. (c) Binned summary heatmap of mean methylation at each of the 23 eligible CpGs, partitioned into six quantile bins of single-read predicted age. Color scale: blue = unmethylated (0.0), red = fully methylated (1.0). (d) Read×CpG binary methylation heatmap for all reads from 20 held-out samples overlapping the window, sorted by single-read predicted age (bottom = youngest; top = oldest). Blue = unmethylated (T / 0); red = methylated (C / 1); white = not covered. Dashed horizontal lines separate Young, Mid, and Old predicted-age groups. The dotted vertical line marks the position of cg16867657 (chr6:11,044,644).

Using reads exclusively from the *ELOVL2* window, sample-level weighted-median predicted age correlated positively with chronological age across the 20 held-out ONT samples (Pearson *r* = 0.64, MAE = 13.5 years; Figure 5b). The regression line broadly tracked the identity line, though with substantial scatter reflecting the inherent noisiness of predictions derived from a single 10 kb window covering only 23 CpGs per read. Despite this, the correlation was detectable and statistically meaningful, indicating that the *ELOVL2* cluster carries genuine age-informative signal at single-molecule resolution.

The binned summary heatmap (Figure 5c) revealed a clear methylation gradient across the *ELOVL2* locus as a function of single-read predicted age. Reads assigned to the youngest predicted-age bin (18.3–32.5 years) showed predominantly unmethylated states across all 23 eligible CpGs, whereas the highest-predicted-age bin (85.3–88.2 years) showed near-complete methylation at the central high-information cluster (chr6:11,044,628–11,044,661), which includes the position corresponding to the canonical array probe cg16867657. Flanking CpGs at the window edges showed weaker age-associated changes, consistent with their lower Fisher information scores.

The read×CpG heatmap sorted by single-read predicted age (Figure 5d) confirmed both the biological signal and a key technical limitation of single-molecule inference at a sparse locus. Reads in the Young group showed predominantly unmethylated states at the central high-information CpGs, while reads in the Old group showed extensive methylation at the same sites. However, a substantial fraction of reads fell near the boundaries of the age grid, visible as dense bands at the top and bottom of the heatmap. This bounda ry-clustering arises because reads covering only 20–23 CpGs provide limited resolution within the interior of the likelihood surface, pushing predictions towards the grid extremes when the methylation pattern is either very low or very high. Sample-level aggregation over dozens of reads mitigates this effect and recovers the expected age correlation, as demonstrated in Figure 5b.

## Discussion

In this study, we developed and evaluated LongReadClock, a set of cell-type-resolved epigenetic aging clocks for blood constructed directly from cohort-scale Oxford Nanopore sequencing data. Applied to the All of Us Research Program, LongReadClock produced accurate bulk and cell-type-specific age predictions across seven blood cell populations, revealed biologically coherent phenotype–EAD associations that were not detected in bulk blood, and demonstrated that cell-type-specific methylation aging signals can be recovered from reads digitally sorted by their methylation patterns alone.

### Lineage specificity validates cell-type clock resolution

The strongest cell-type-specific association observed in this study involved sex and lymphoid EAD. Male sex was associated with lower B cell and NK cell EAD, whereas no corresponding association was observed in bulk blood or the myeloid-lineage clocks. This pattern is broadly consistent with established sexual dimorphism in lymphoid immune regulation. Females generally exhibit stronger humoral immune responses, while sex hormones directly modulate B cell survival and activation and regulate NK cell responses [30, 31, 32]. These differences provide a potential biological context for the lower B cell and NK cell EAD observed in males, although the mechanisms linking sex-dependent immune regulation to cell-type-specific epigenetic aging remain unknown.

Several other observed associations were also broadly consistent with established lineage biology. The associations of systolic blood pressure and resting heart rate with myeloid-lineage EAD are consistent with the established involvement of monocytes and neutrophils in cardiovascular inflammation and injury [33]. The association between BMI and B cell EAD is similarly consistent with experimental evidence implicating B cells in obesity-associated inflammation and metabolic dysfunction [34]. These associations suggest that cell-type-resolved EAD can reveal association patterns that are not apparent in bulk blood.

### Comparison with prior approaches

LongReadClock extends the LongReadAge framework [24] to cohort scale and to a broader set of cell-type populations, and, to our knowledge, provides the first systematic evaluation of cell-type-resolved epigenetic clocks in a large long-read sequencing dataset. Prior cell-type-specific clocks have been developed on methylation arrays [20], which require physical cell sorting and are limited by the fixed probe coverage of array platforms. In contrast, LongReadClock operates on digitally sorted reads from bulk blood samples, requires no additional assays or physical separation, and is applicable to any sample for which long-read methylation data are available.

Statistical deconvolution methods adjust bulk methylation profiles for estimated cell-type proportions post hoc [18, 19]. LongReadClock operates at a fundamentally different level of resolution: rather than adjusting a bulk signal, it classifies individual sequencing reads with target-specific cell-type models before age inference, producing cell-type-enriched methylation profiles without estimating cell proportions from bulk averages. This read-level approach is only possible because long reads span multiple CpGs in a contiguous haplotype, enabling methylation pattern matching at the molecular rather than population level.

Elastic net regularised regression achieved slightly lower bulk blood MAE than the likelihood - based approach on the held-out test set, consistent with prior observations that global penalised regression can outperform site-by-site likelihood models for bulk tissue [11, 12]. However, the elastic net is not applicable to cell-type-specific inference, where CpG observations per read are sparse and a feature matrix cannot be constructed. The likelihood-based approach therefore provides practical path to cell-type-resolved age inference from long-read data at present.

### Single-molecule age inference at the ELOVL2 locus

The exploratory single-molecule analyses demonstrated that individual long reads carry detectable age-informative methylation signal, with sample-level aggregation over the ELOVL2 window yielding a Pearson correlation of 0.64 with chronological age. This supports the feasibility of the LongReadAge approach [24] at a locus with exceptionally high per-CpG informativeness. However, the boundary clustering of single-read predictions illustrated a fundamental information-theoretic constraint: individual reads covering tens of CpG sites carry insufficient information to resolve age within the interior of the likelihood surface, and most predictions collapse to the age grid boundaries. These results show that accurate single - molecule epigenetic age inference was not achieved from this 10-kb window. Longer reads or targeted constructs spanning many more informative CpGs may be needed to overcome this constraint

### Methylation call quality and future cohort expansion

The present analysis focused on ONT data because platform-specific differences in methylation-call confidence substantially affected age inference. Filtering intermediate-confidence methylation calls improved the performance of PacBio-derived profiles but reduced effective CpG coverage, and models incorporating PacBio data did not outperform those trained on the smaller, higher-confidence ONT cohort. These observations suggest that increasing cohort size does not necessarily improve methylation-based age inference when additional samples contribute lower-quality or substantially sparser methylation information.

The differences observed here may reflect the methylation calls available under the basecalling models used in the All of Us v8 release rather than a fundamental limitation of PacBio sequencing chemistry. Because LongReadClock operates on called methylation states rather than platform-specific raw signal, improvements in methylation calling could enable future integration of PacBio data and substantially expand the available cohort.

The All of Us Research Program is continuing to expand its long-read sequencing cohort, and future data releases may provide substantially larger numbers of samples suitable for cell-type-resolved epigenetic age inference. Such expansion would increase statistical power and enable more rigorous evaluation and replication of the cell-type-specific associations observed here. Longitudinal data could further establish whether cell-type-specific EAD predicts incident disease or other future health outcomes.

Beyond All of Us, the analytical framework underlying LongReadClock could be adapted to other long-read methylation datasets and potentially to tissues beyond blood. Application to other tissues would require appropriate cell-type methylation references and tissue-specific age models rather than direct transfer of the blood-trained clocks. In tissues with heterogeneous cell composition, however, the combination of methylation-based digital read sorting and sparse-observation age inference may provide a general strategy for resolving cell-type-specific epigenetic aging without physical cell sorting. Single-cell or spatially resolved long-read methylation data, for which sparse-observation likelihood approaches are particularly well suited [23], could extend this framework to more highly resolved biological settings. Training cell-type-specific reference models on physically sorted long-read data, and incorporating haplotype-level or local methylation patterns observed on individual long reads, could further improve digital sorting and age prediction.

### Limitations

The All of Us long-read whole-genome sequencing dataset is currently cross-sectional, so the phenotype associations reported here cannot establish causal directionality or determine whether cell-type-specific EAD predicts future health outcomes. Longitudinal studies will be required to evaluate the prognostic value of LongReadClock.

Phenotype association analyses were restricted to the 324 ONT participants with high-quality methylation calls, limiting statistical power to detect modest associations and necessitating cautious interpretation of both nominal associations and null findings. The primary ONT cohort was also predominantly self-reported Hispanic/Latino, and the generalisability of the observed associations to other populations remains unknown. Replication in larger and ancestrally diverse cohorts will therefore be important.

Digital read sorting was evaluated against purified-cell reference methylation profiles, but direct validation against long-read data from physically sorted cell populations was not available, and the effect of read-assignment stringency was not systematically assessed.

Coverage represents an additional constraint on cell-type-specific inference. Because target-specific sorting retains only a subset of reads for each output, lower-abundance cell types receive fewer reads and consequently have sparser CpG coverage. This limitation was most apparent for the B cell and NK cell clocks, which showed lower prediction accuracy than the more abundant cell populations. Greater sequencing depth and larger cohorts may therefore be particularly important for resolving epigenetic aging in low-abundance cell types.

The likelihood model treats CpG methylation states as conditionally independent given age, although neighbouring CpGs can exhibit substantial co-methylation. In addition, the grid-based age estimator can produce boundary effects when methylation information is sparse. This was most apparent in the single-molecule ELOVL2 analysis, where individual reads covered only 20–23 informative CpGs and frequently yielded predictions near the age-grid boundaries. Inference methods that more explicitly model uncertainty and dependencies among CpGs may improve performance, particularly in sparse single-read settings.

## Conclusion

LongReadClock demonstrates that cell-type-resolved epigenetic age inference is feasible at cohort scale from standard bulk long-read blood methylation data, in the absence of cell sorting. The digital sorting pipeline recovers lineage-enriched aging signals not apparent in bulk analyses, produces accurate age predictions across seven blood cell populations, and identifies biologically coherent phenotype associations.

These results support long-read methylation sequencing as a practical substrate for cell-type-resolved epigenetic aging research and provide a foundation for larger studies linking cell-type-specific epigenetic aging to human health and disease.

## Materials and Methods

### Long-read whole-genome sequencing data and methylation preprocessing

#### Data source

Long-read whole-genome sequencing (lrWGS) data were obtained from the All of Us Research Program Controlled Tier CDR release C2024Q3R9 (v8) [27]. The available data comprised 2,842 platform-specific sequencing records generated across multiple facilities and sequencing platforms. Because some participants were sequenced on more than one platform, these record counts do not represent 2,842 unique individuals.

Oxford Nanopore Technology R10.4 data were generated on PromethION instruments at Baylor College of Medicine (BCM; *n* = 196, high-pass 25×) and Johns Hopkins University (JHU; *n* = 128, high-pass 25×). PacBio Revio data were generated at BCM (*n* = 111, high-pass 25×), the Broad Institute (BI; *n* = 803, mid-pass 12×), and the University of Washington (UW; *n* = 53, high-pass 25×). PacBio Sequel IIe data were generated at BCM (*n* = 77, high-pass 25×), BI (*n* = 84 high-pass 25× and *n* = 198 mid-pass 12×), and UW (*n* = 100, high-pass 25×). The dataset also included legacy PacBio Sequel IIe and Sequel II records from the Heliogen cohort available in the CDRv7 Controlled Tier (*n* = 1,027, mid-pass 12×), together with an additional Heliogen PacBio Revio subset (*n* = 65, mid-pass 12×).

Following platform-specific quality assessment (see Methylation calling quality control), the 324 ONT records from BCM and JHU were selected as the primary analytic cohort for LongReadClock reference construction, internal cross-validation, and the primary phenotype association analyses. For cross-platform validation, 459 PacBio Sequel IIe and 1,032 PacBio Revio records were additionally processed using the high-confidence methylation-call filter described below. The resulting ONT and PacBio analysis set comprised 1,815 platform-specific records from 1,773 unique participants.

Platform-specific reference models were constructed from the ONT, Sequel IIe, Revio, and combined training cohorts for cross-platform evaluation. The legacy CDRv7 records (C2022Q4R9) were retained for platform-level quality assessment but were excluded from clock construction and downstream phenotype analyses because their available methylation representations did not retain informative variation at the evaluated age-associated loci.

#### Methylation calling and file conversion

CpG methylation information was extracted from modification-aware BAM files using `wgbstools` [35]. Base-modification calls and their associated probabilities were obtained from the MM and ML auxiliary tags generated by the platform-specific methylation-calling workflows. ML values were encoded on an integer scale from 0 to 255.

For PacBio samples, calls with ML scores from 51 to 204, corresponding approximately to modification probabilities from 0.20 to 0.80, were excluded before downstream processing. Calls outside this interval were retained as high-confidence unmethylated or methylated observations. The resulting high-confidence PacBio representations were used for reference construction and cross-platform evaluation. ONT samples were processed without this additional probability filter.

Two methylation representations were generated for each sample. The `.pat.gz` format stores BGZF-compressed, tab-separated read-level methylation patterns. Each record contains the chromosome, the 1-based genome-wide CpG index of the first represented CpG, a methylation string spanning consecutive CpGs (`C` = methylated, `T` = unmethylated, and . = no retained methylation observation), and the number of reads sharing the same genomic start and methylation pattern. The `.beta` format stores a compact binary array indexed by genome-wide CpG position. Each CpG is represented by paired values corresponding to methylated-read count and total read coverage.

The hg38 reference genome was initialized using `wgbstools init_genome`, which generated a genome-wide CpG coordinate index shared across samples. BAM files were converted using `wgbstools bam2pat`; with aggregate output enabled, the conversion generated both read-level `.pat.gz` files and per-CpG `.beta` files. All downstream methylation profiles were indexed in this common 1-based CpG address space.

#### Computational infrastructure

ONT BAM-to-PAT conversion was parallelised across samples using `dsub` [36] with the Google Cloud Batch backend in the All of Us Researcher Workbench. Each sample was processed as an independent task on a preemptible Google Cloud virtual machine using a Cloud SDK container.

Python wheels, `wgbstools`, and the hg38 reference assets were pre-staged in workspace-controlled storage and installed or unpacked locally at task startup. Each task initialised the hg38 `wgbstools` reference, retrieved the sample BAM and corresponding index from controlled-tier requester-pays storage, executed `wgbstools bam2pat`, and wrote the resulting `.pat.gz` and `.beta` files to project storage. Generated files were accessed through the mounted workspace storage for subsequent analyses.

### Methylation calling quality control

#### Modification-tag interpretation

Base modifications were represented by the paired `MM` and `ML` SAM auxiliary tags. The `MM` tag identifies the modification type and its positions along the read, while the corresponding `ML` byte array stores a linearly scaled probability for each listed modification call. The evaluated calls represented CpG 5-methylcytosine (5mC). Low `ML` values therefore indicated low support for 5mC, whereas high values indicated strong support for 5mC.

`ML` values ranged from 0 to 255. Values from 51 to 204 corresponded approximately to modification probabilities between 0.20 and 0.80 and were classified as intermediate - confidence calls. Values outside this range were classified as high-confidence unmethylated or methylated calls.

#### Principal component analysis of bulk methylation profiles

Global technical structure was assessed using bulk methylation profiles from the 2,820 samples with available `.beta` files. A reproducible random sample of 50,000 CpG indices was selected from the hg38 CpG index (random seed 20260426). Samples were required to have observations at at least 50% of sampled sites, and CpGs were retained when observed in at least 80% of samples. Missing values at retained sites were imputed using the site-wise mean. Randomised principal component analysis was then performed using scikit-learn [37], and principal-component scores were visualised by sequencing platform and source batch.

#### ML-score ambiguity metric

For each v8 ONT, PacBio Revio, and PacBio Sequel IIe sample, the first 50,000 mapped alignments were streamed from the modification-aware BAM file. The `ML` byte arrays were extracted from alignments containing modification tags.

The ambiguous fraction was defined as

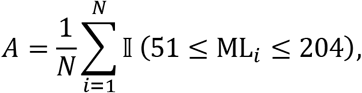

where *N* is the total number of extracted `ML` entries and I is the indicator function. Higher values indicated a greater proportion of intermediate-confidence modification calls.

#### Implementation

BAM data were streamed without writing complete alignment files to local storage using `gcloud storage cat` piped to `samtools view`. Per-process timeouts were used to terminate stalled streams. Each sample was processed independently, and sample-level QC results were serialised to JSON and written incrementally. The procedure was applied to 1,815 v8 samples: 324 ONT, 1,032 PacBio Revio, and 459 PacBio Sequel IIe samples.

#### Platform comparison and sample selection

Ambiguous-fraction distributions were summarised by sequencing platform and source batch (Figure 2b–c). The 324 ONT samples from BCM and JHU were selected as the primary cohort for LongReadClock construction and phenotype association analyses.

For cross-platform analyses, PacBio modification calls with `ML` values from 51 to 204 were excluded before generation of the filtered `.pat.gz` and `.beta` representations. The existing ONT representations were retained unchanged. Platform-specific ONT, PacBio Sequel IIe, PacBio Revio, and combined references were constructed from these data for cross-platform evaluation. The ONT-trained references were retained as the primary LongReadClock models.

Legacy Controlled Tier v7 samples were excluded from clock construction after their bulk methylation profiles showed negligible informative methylation variation.

### Digital cell-type sorting of long reads

#### Overview

Long-read methylation patterns were classified using target-specific likelihood-ratio models adapted from [24]. Seven reference models were evaluated: myeloid, lymphoid, T cell, B cell, NK cell, monocyte, and granulocyte. The myeloid and lymphoid models used reciprocal lineage backgrounds, whereas the five subtype models used one-versus-all reference backgrounds.

#### Read classification by likelihood-ratio scoring

Each PAT pattern record was evaluated at informative CpGs overlapping the marker regions for target model *k*. A PAT record represents one or more reads sharing the same genomic start position and methylation pattern. Let *m_i_* = 1 denote methylation and *m_i_* = 0 denote unmethylation at CpG *i*. The target-versus-background log-likelihood ratio was

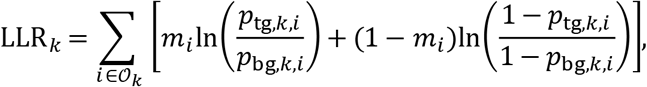

where *p*_tg,*k*,*i*_ and *p*_bg,*k*,*i*_ are the target and background methylation fractions encoded in the corresponding marker BED file. Probabilities were clipped to [10^−4^, 1 − 10^−4^] to avoid undefined logarithms. Unknown or ambiguous PAT states were omitted from the score.

A PAT record was retained in target-specific output *k* when it covered at least five informative CpGs and LLR*_k_* > ln(1.053). Records with LLR*_k_* < −ln(1.053) were recorded as background-favouring for that model, whereas records with intermediate scores or fewer than five informative CpGs were excluded from that target-specific output. The seven models were evaluated independently in a single pass through each PAT file. Consequently, a record satisfying more than one target model could contribute to multiple target-specific outputs.

#### Output file generation

Records retained by each target model were written to target-specific PAT files. These files were compressed, indexed, and converted to beta format using `wgbstools pat2beta` [35]. The beta files contained methylated and total read counts at each CpG in the hg38 CpG coordinate index and were used for downstream reference construction and age inference.

#### Computational infrastructure

Cell-type classification was parallelised by sample using `dsub` [36] and Google Cloud Batch. Each job loaded the seven marker models, classified one PAT file, converted nonempty target-specific outputs to beta format, and wrote the resulting PAT, index, beta, and classification-summary files to workspace storage.

#### Validation against purified-cell profiles

Digital sorting was evaluated by comparing ONT-derived myeloid and lymphoid methylation profiles with purified-cell WGBS profiles from the Human Cell Type DNA Methylation Atlas [29]. Genomic overlap intervals between the reciprocal myeloid and lymphoid marker sets were identified, and intervals shorter than 50 bp were removed. One hundred eligible intervals were sampled using a fixed random seed.

Purified myeloid profiles were constructed by averaging sample-level methylation fractions across the monocyte and granulocyte reference samples. Purified lymphoid profiles were similarly constructed from the T cell, B cell, and NK cell reference samples. Digitally sorted myeloid and lymphoid profiles were constructed by averaging sample-level methylation fractions across 65 ONT participants with both target-specific outputs.

For each interval containing at least five CpGs observed in all four aggregate profiles, mean absolute methylation differences were calculated for lineage-matched comparisons (digitally sorted myeloid versus purified myeloid and digitally sorted lymphoid versus purified lymphoid) and lineage-mismatched comparisons (digitally sorted myeloid versus purified lymphoid and digitally sorted lymphoid versus purified myeloid). Concordance was evaluated by comparing the distributions of mean absolute methylation differences between matched and mismatched profile pairs.

### Age reference model construction and likelihood-based age inference

#### Age reference model

For training participant *j*, the methylation fraction at CpG *i* was calculated as *y_ij_* = *m_ij_*/*c_ij_* when total coverage *c_ij_* > 0, where *m_ij_* and *c_ij_* denote methylated and total read counts, respectively. Per-CpG associations with age were estimated by unweighted ordinary least-squares regression of methylation fraction on chronological age. Thus, each participant with nonzero coverage contributed one sample-level methylation fraction, irrespective of sequencing depth.

Sufficient statistics were accumulated incrementally across training samples to avoid loading all beta files simultaneously. These statistics were used to calculate the slope γ̂*_i_*, intercept α̂*_i_*, Pearson correlation *r_i_*, and correlation *t*-statistic

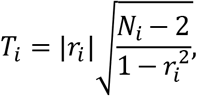

where *N_i_* is the number of training participants with nonzero coverage at CpG *i*. The absolute *t*-statistic *T_i_* was used to rank age-informative CpGs.

#### Eligible site selection

A CpG was retained when it was observed in at least

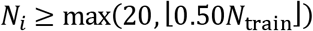

training participants and had |*r_i_*| ≥ 0.30. The endpoints of the training-derived 0.1-year age grid were defined as

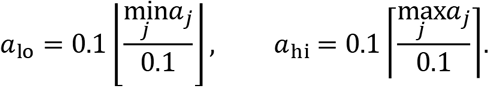

Predicted methylation fractions at both endpoints were required to lie strictly within (10^−3^, 1 − 10^−3^). The absolute predicted methylation change across these endpoints was required to satisfy

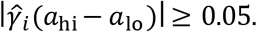

Sites passing all criteria formed the eligible CpG set for each reference model.

#### Likelihood-based age inference

For each new sample, covered eligible CpGs were ranked by their training-set statistic *T_i_*, and up to the top *K* = 1,000 sites were retained. Predictions required at least 10 covered eligible sites.

To limit the influence of unusually high coverage, total depth at each selected CpG was capped at 20:

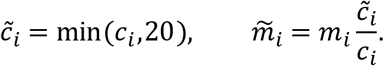

This transformation preserved the observed methylation fraction while limiting the effective contribution of each CpG. At candidate age *a*, the reference probability was

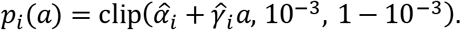

Age was estimated using the depth-capped, binomial-form weighted log-likelihood

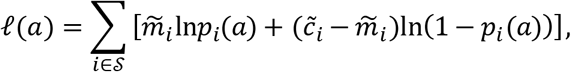

where *S* is the selected set of covered eligible CpGs. The predicted age was

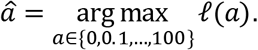

#### Selection of the number of informative CpGs

The number of CpGs retained for inference was evaluated using *K* ∈ {50,100,200,500,1000,2000}. For each value of *K*, age was inferred in the held-out ONT test set using the same autosomal 0–100-year grid. Performance was summarized by Pearson correlation, mean absolute error, mean bias, predicted-age standard deviation, and the median number of CpGs used per prediction. Performance was stable near *K* = 1,000, which was selected for the primary analyses.

#### Elastic-net comparison

For comparison with a conventional regression approach, an elastic-net model was trained on bulk ONT methylation fractions at the same top 1,000 informative autosomal CpGs. CpGs observed in fewer than 20% of training participants were removed. Remaining missing values were imputed using training-set CpG means, and features were standardized using training-set means and standard deviations.

The mixing parameter was evaluated over *l*_1__ratio ∈ {0.1,0.5,0.9,1.0}, and the regularization parameter *α* was evaluated at 30 logarithmically spaced values between 10^−3^ and 10^1^. Hyperparameters were selected by five-fold cross-validation within the training set. The selected model was then fitted to the complete training set and evaluated once on the held-out ONT test participants. No held-out test information was used for feature filtering, imputation, scaling, or model selection.

#### Five-fold cross-validation

Out-of-fold predictions for downstream epigenetic age deviation analyses were generated by five-fold participant-level cross-validation within the ONT cohort. Reference fitting, eligible-site selection, and age inference were repeated independently within each training fold. Each participant therefore received a prediction from a reference model that had not been trained on that participant. These out-of-fold predictions were used to calculate epigenetic age deviation as described in Methods, Phenotype association analysis.

#### Cross-platform validation

Cross-platform validation used PacBio methylation representations in which calls with modification probabilities in the interval [0.20,0.80] were omitted before PAT and beta generation. Reference models were constructed separately from ONT, PacBio Sequel IIe, PacBio Revio, and pooled long-read training samples.

Each reference was evaluated on held-out ONT, Sequel IIe, Revio, and pooled test manifests for bulk blood and all seven digitally sorted compartments. Age inference used *K* = 1,000 CpGs and the fixed 0–100-year grid, producing 128 reference-by-test-platform-by-compartment evaluations. Matched-platform evaluations were held out from their corresponding reference training sets; the same fixed platform-specific test manifests were used for cross-platform comparisons. Performance was summarized using mean absolute error and Pearson correlation between chronological and predicted age.

### Phenotype association analysis

#### Epigenetic age deviation

Epigenetic age deviation (EAD) was defined as predicted age minus chronological age. Primary ONT estimates were obtained by five-fold cross-validation, such that each participant was evaluated using references trained on the other four folds. Out-of-fold EAD was calculated for bulk blood and the seven cell-type-specific clocks.

For the descriptive comparisons in Figure 4a–b, age-adjusted EAD residuals from the canonical 324-participant ONT dataset were used. Pearson correlations were calculated between clocks, and the proportion of cell-type EAD variance not explained by bulk EAD was summarized as 100(1 − *r*^2^).

#### Phenotype definitions

Phenotypes were extracted from the All of Us Controlled Tier C2024Q3R9 dataset. Sex at birth was coded as male or female, with other responses treated as missing. Continuous measurements were restricted to prespecified biologically plausible ranges and averaged across qualifying records within each participant.

BMI, systolic blood pressure, and resting heart rate were extracted using concepts 3038553, 903118, and 903126, respectively. HbA1c combined two equivalent standard concepts (3004410 and 3005673). eGFR was defined using the race-unspecified MDRD estimate (LOINC 77147-7; OMOP concept 46236952), selected as the non-race-stratified eGFR concept with the greatest participant coverage. Values were restricted to 5–200 mL/min/1.73 m^2^.

Clinical conditions were defined from qualifying records in the OMOP condition occurrence table of the All of Us CDR. Obesity was defined as either an obesity diagnosis or mean BMI ≥ 30 kg/m^2^. Medication exposures were mapped to ingredient concepts using the OMOP concept ancestor hierarchy. Lifestyle and self-reported health variables were derived from All of Us survey responses; current smokers included participants reporting at least 100 lifetime cigarettes and current daily or occasional smoking, with former smokers excluded from that comparison.

The complete exploratory screen included 31 demographic, lifestyle, clinical, medication, laboratory, and health burden phenotypes. Figure 4 presents a focused subset, and complete results are reported in Supplementary Table S1.

#### Association models

Each phenotype was tested independently against each clock using complete-case ordinary least squares regression. Models required at least 30 participants and included linear and quadratic chronological age. Non-sex phenotypes were additionally adjusted for sex. For bulk, myeloid, monocyte, and granulocyte clocks, a sex-by-age interaction was included based on model comparisons:

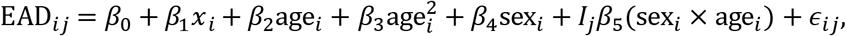

where *I_j_* = 1 for clocks containing the interaction and zero otherwise. When sex was the phenotype, it was treated as the primary predictor. For clocks containing the interaction, the male–female contrast was evaluated at the mean age of the complete-case sample.

Regression coefficients represent the associated difference in mean EAD per unit increase in a continuous phenotype or between groups for a binary phenotype. Conventional OLS standard errors, two-sided *p*-values, and 95% confidence intervals were reported.

### Single-molecule age inference

#### Fisher information–guided window selection

To identify genomic regions with high potential information for single-molecule age inference, we calculated Fisher information at each eligible CpG in the bulk ONT reference model, which was fitted using the ONT training set. For a site with fitted methylation probability *p_i_*(*a*) = *β*_0,*i*_ + *β*_1,*i*_*a*, Fisher information for age was calculated as

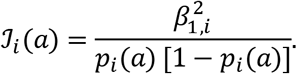

Predicted probabilities were clipped to the numerical bounds used during reference construction before calculating Fisher information. Site-level Fisher information was averaged over a uniform age grid from 20 to 100 years in one-year steps.

The genome was tiled into overlapping 10-kb windows at 2.5-kb steps, and mean site-level Fisher information was summed across eligible CpGs within each window. Windows were ranked by summed Fisher information. The *ELOVL2*-centred region was among the most informative regions identified by this screen and was selected for detailed analysis because of the established relationship between *ELOVL2* methylation and chronological age. The evaluated region spanned chr6:11,039,644–11,049,644 and contained 23 eligible CpGs.

#### Single-read age scoring

Single-molecule age inference was evaluated in 65 held-out ONT participants. PAT records overlapping the *ELOVL2* region were parsed to recover methylation states at eligible CpGs. Each PAT record contains a methylation pattern over consecutive CpGs (`C` = methylated, `T` = unmethylated, and . = unobserved or ambiguous) and a multiplicity indicating the number of reads represented by that pattern.

Records covering at least three eligible CpGs were retained. For record *r*, age was inferred by maximizing

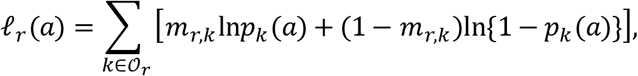

where O*_r_* denotes the eligible CpGs observed in the record, *m_r_*_,*k*_ ∈ {0,1} is the observed methylation state, and *p_k_*(*a*) = *β*_0,*k*_ + *β*_1,*k*_*a* is the fitted reference probability. Probabilities were clipped to the numerical bounds used during reference construction. The likelihood was evaluated over an age grid from 0 to 100 years in 0.1-year increments.

A sample-level *ELOVL2* age estimate was calculated as the weighted median of its pattern-level age estimates, using PAT multiplicity as the weight.

#### Read-pattern-by-CpG methylation visualisation

For visualisation, a methylation matrix was constructed with one row per retained PAT pattern record and one column per eligible CpG in genomic order. Matrix entries represented methylated states (1), unmethylated states (0), or missing coverage. Rows were ordered by their inferred single-molecule age.

For the full pattern-level heatmap, the 10th and 90th percentiles of the inferred-age distribution were used to mark young-boundary, middle, and old-boundary groups. A summary heatmap was constructed using fixed 10-year predicted-age intervals from 0 to 100 years. Within each interval, mean methylation was calculated at each eligible CpG using the observed pattern records. PAT multiplicities were used for sample-level age summaries but were not used to duplicate rows in either heatmap.

## Acknowledgements

This work was supported by the National Institute on Aging and Hevolution Foundation grants to VNG. We gratefully acknowledge the National Institutes of Health All of Us Research Program for providing access to omics data and participant information.

## Author Contribution Statement

YW and VNG conceived and designed the study. YW curated the data, implemented the code, performed the analyses, and wrote the manuscript. AE provided methodological guidance. HS, MM, and VNG reviewed and edited the manuscript. VNG supervised the study and acquired funding.

## Data Availability

This study used data from the All of Us Research Program Controlled Tier Dataset v8. Authorized researchers may apply for access through the All of Us Researcher Workbench. Individual-level participant data cannot be distributed by the authors. Analysis code will be made publicly available through GitHub upon publication.

## Ethics

All of Us participants provided informed consent under a protocol reviewed and approved by the All of Us Institutional Review Board. This secondary analysis used Controlled Tier data within the All of Us Researcher Workbench and was conducted in accordance with All of Us data-use policies.

