## supplementary materials for "Measuring Cell Type Resolved Aging Through Long-Read Sequencing"

#### Supplementary Figures

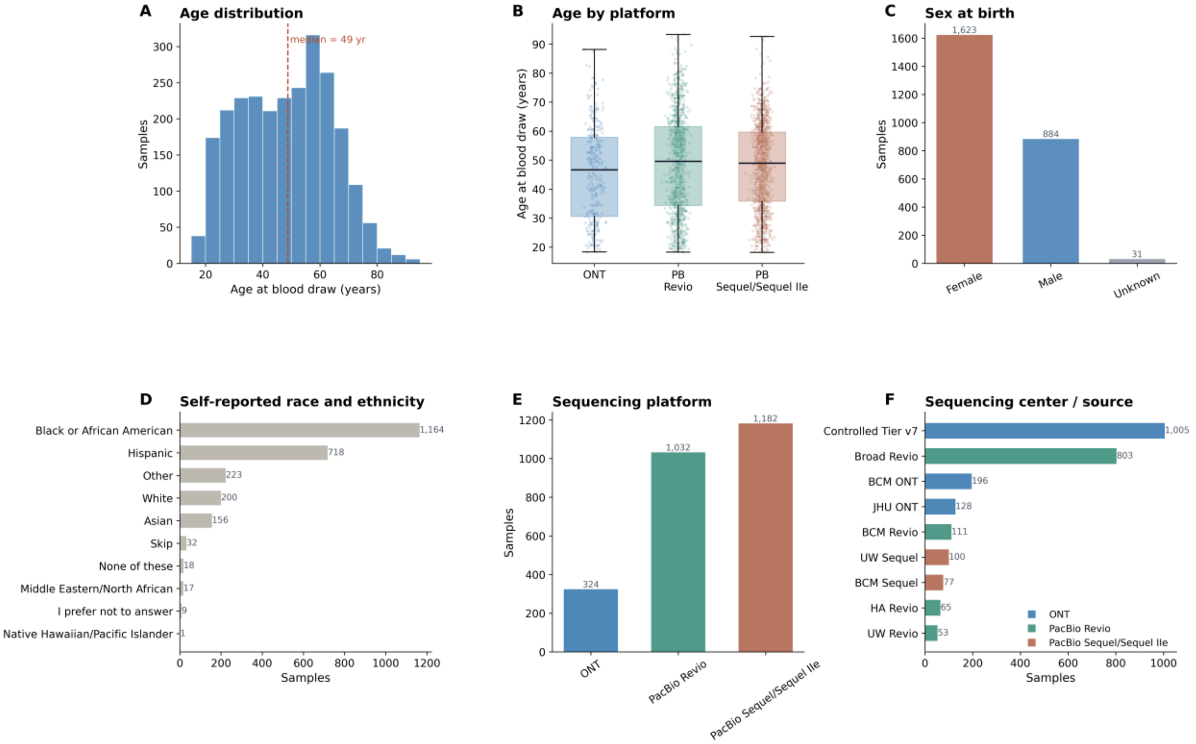


**Supplementary Figure S1.** Cohort composition of the All of Us long-read methylation dataset. Shown are the age distribution, age by platform, sex at birth, self-reported race and ethnicity, sequencing platform, and sequencing center or source.

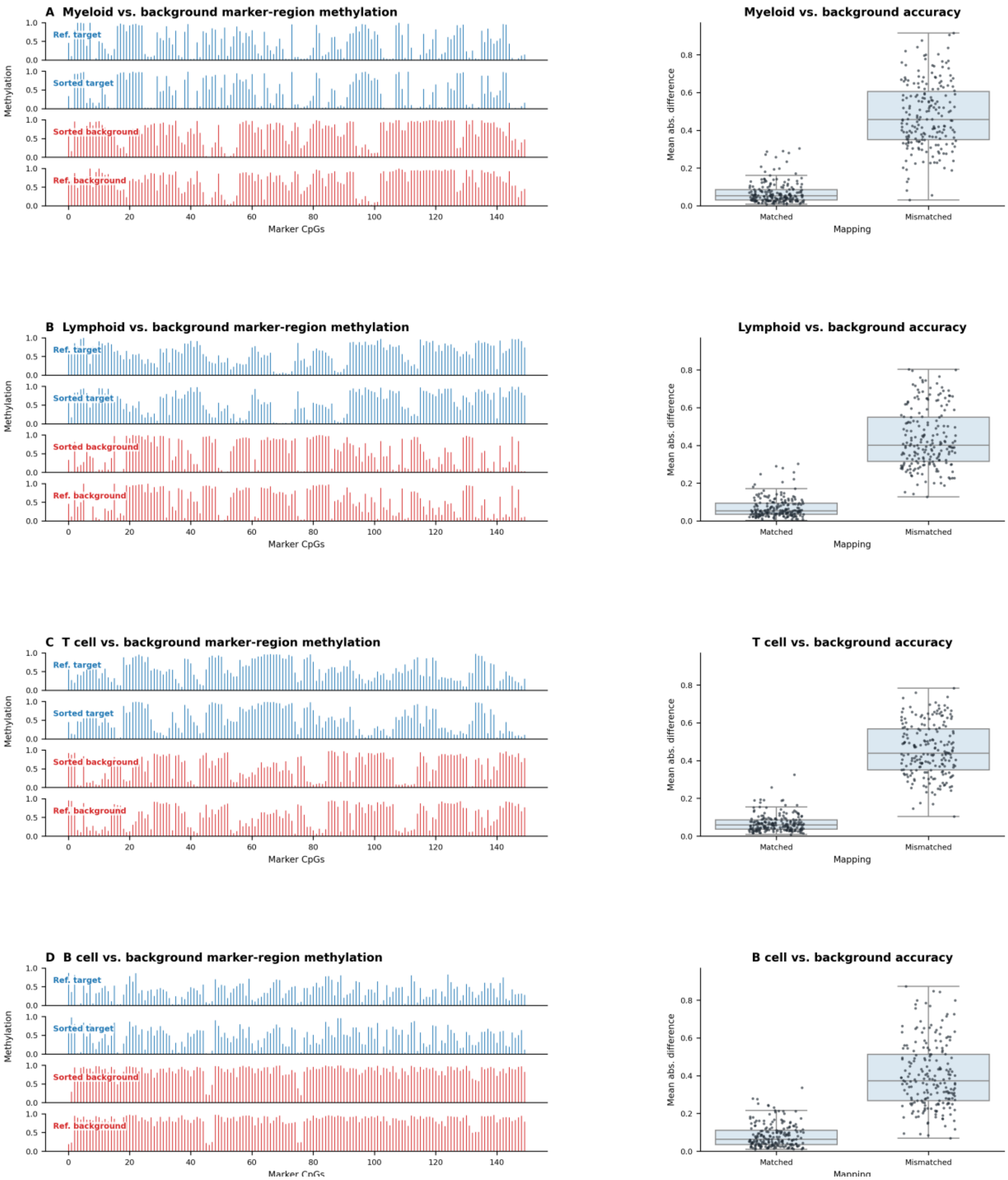


**Supplementary Figure S2.** Validation of target-specific methylation profiles across digitally sorted cell types. (a–d) Myeloid, lymphoid, T cell, and B cell targets, respectively. Left, purified-cell reference and averaged ONT read methylation profiles at target-specific marker CpGs for reads assigned to the target or background. Right, per-marker-region mean absolute methylation differences for matched and mismatched sorted-reference pairs.

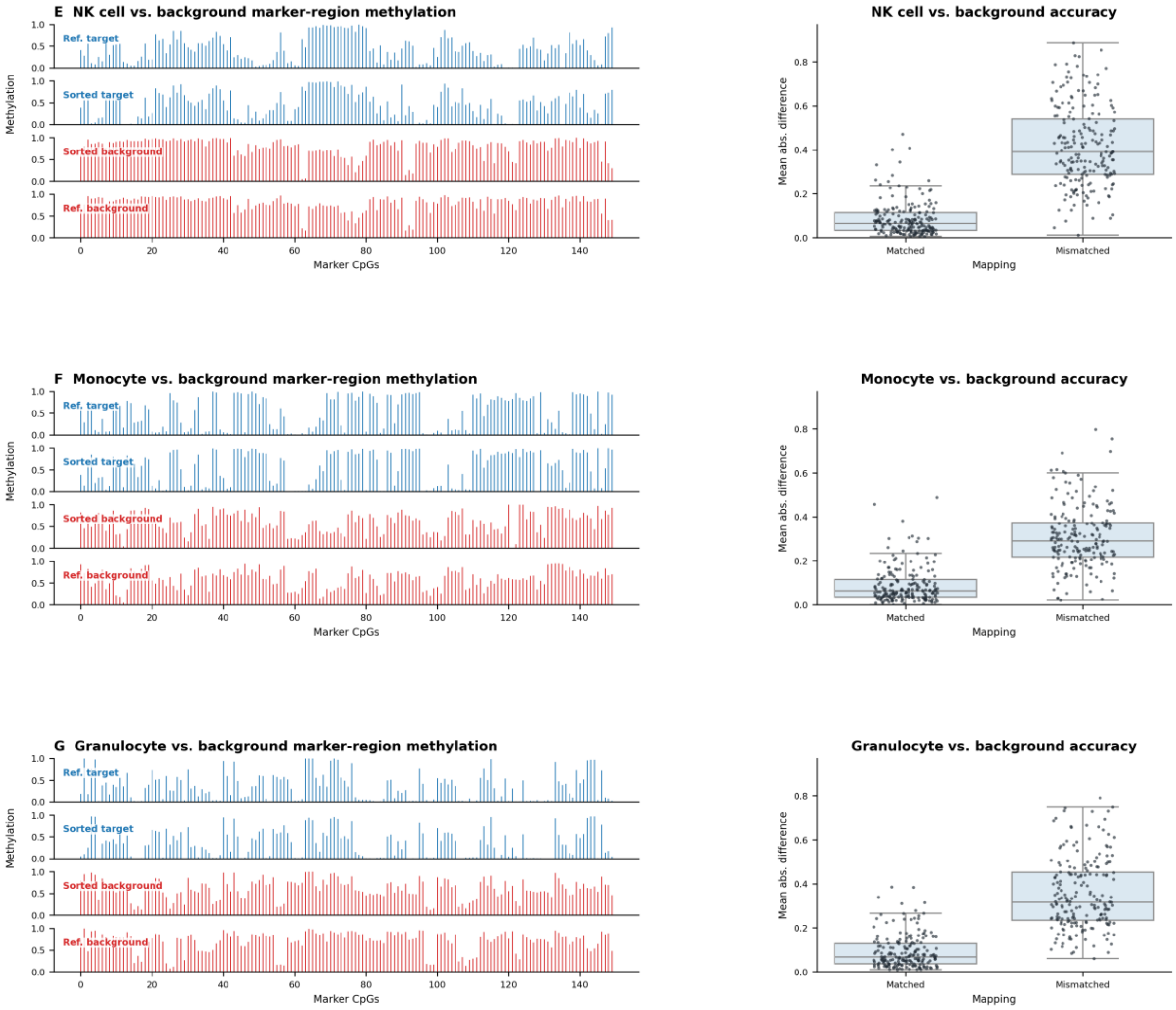


**Supplementary Figure S2 (continued).** Validation of target-specific methylation profiles across digitally sorted cell types. (e–g) NK cell, monocyte, and granulocyte targets, respectively.

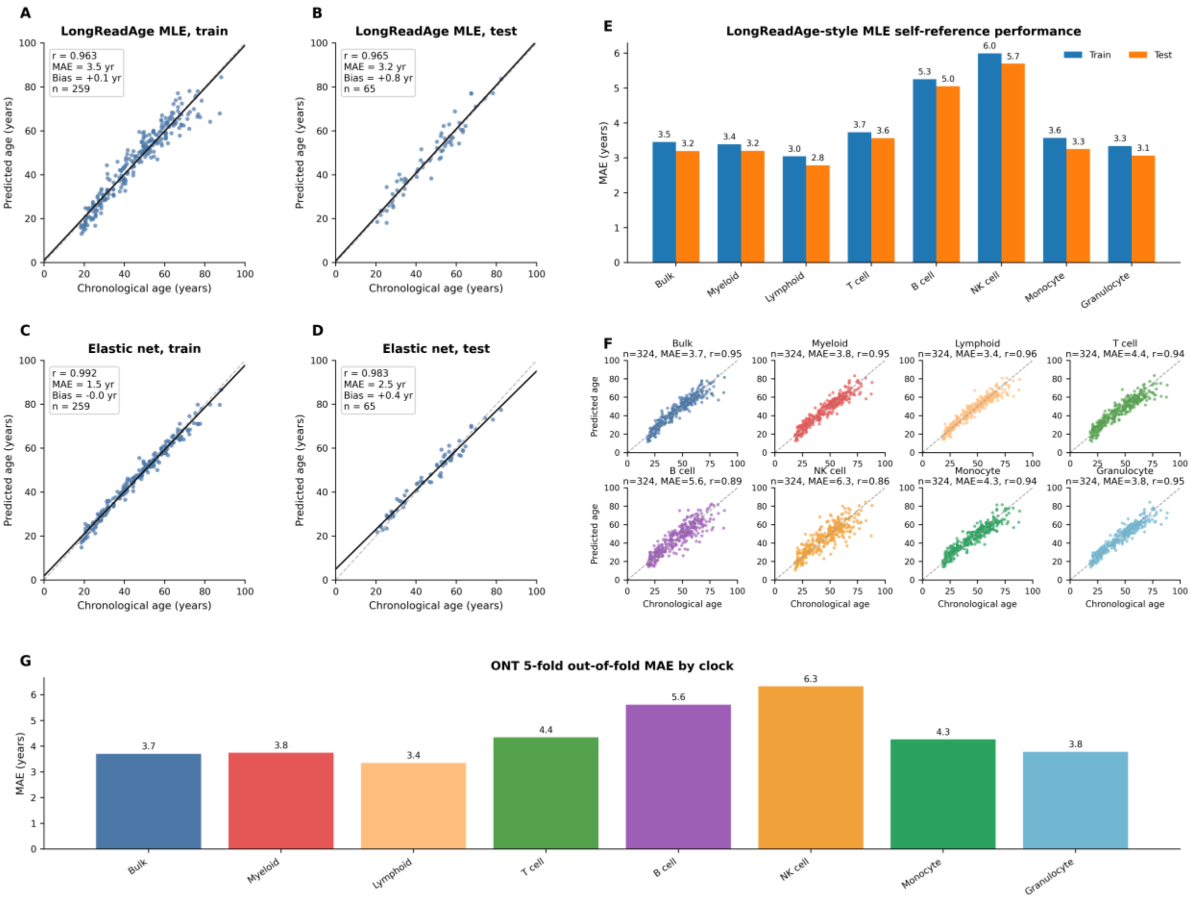


**Supplementary Figure S3.** Comparison of bulk ONT age prediction across likelihood-based and elastic net models, together with train-test and cross-validation performance. Left, bulk ONT age prediction using the LongReadAge-style likelihood-based approach and elastic net, shown separately for training and held-out test sets. Elastic net achieved slightly lower error in bulk prediction. Right, comparison of train and test self-reference performance across bulk and cell-type-specific clocks, five-fold out-of-fold prediction plots, and cross-validated MAE summaries, showing little evidence of overfitting.

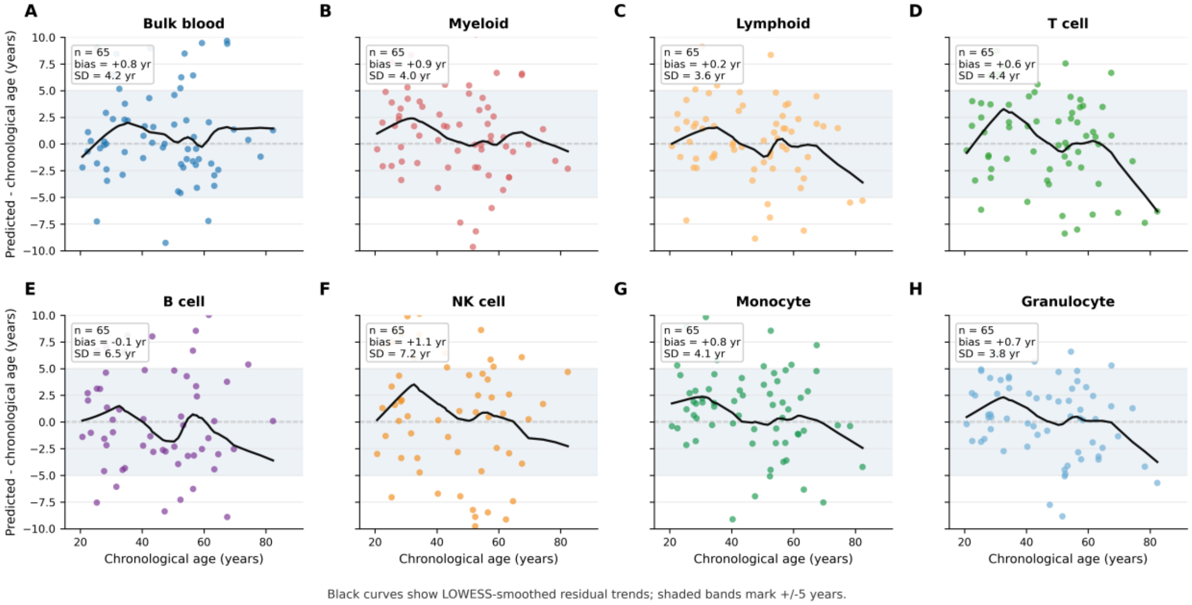


**Supplementary Figure S4.** Residuals versus chronological age for bulk and cell-type-specific ONT self-reference clocks. Residuals are defined as predicted age minus chronological age. Smoothed trends suggest only modest regression-to-the-mean effects overall, with larger apparent deviations at older ages where sample counts are sparse, particularly for some lower-coverage cell types.

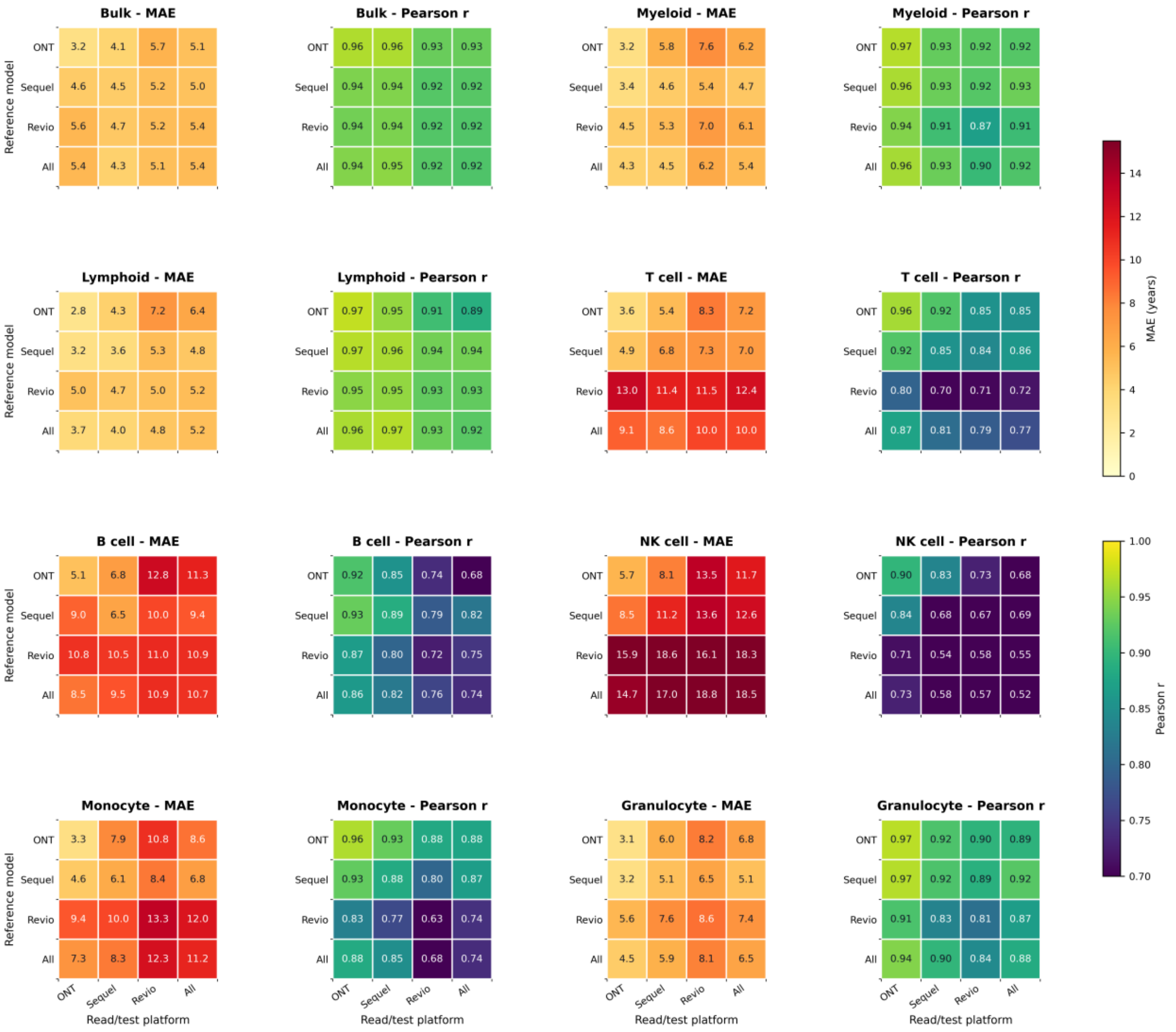


**Supplementary Figure S5.** Cross-platform performance of bulk methylation age prediction using platform-specific reference models. Each heatmap cell summarizes performance when applying the row platform's bulk reference model to the column platform's bulk read data, using \(K=1000\) age-informative CpG sites. Metrics shown are mean absolute error, bias, Pearson correlation, and predicted age standard deviation. Across all evaluated test subsets, including the combined cohort and PacBio-derived test sets, the ONT-trained reference consistently showed the strongest overall performance, indicating that the ONT reference generalized best across platforms.

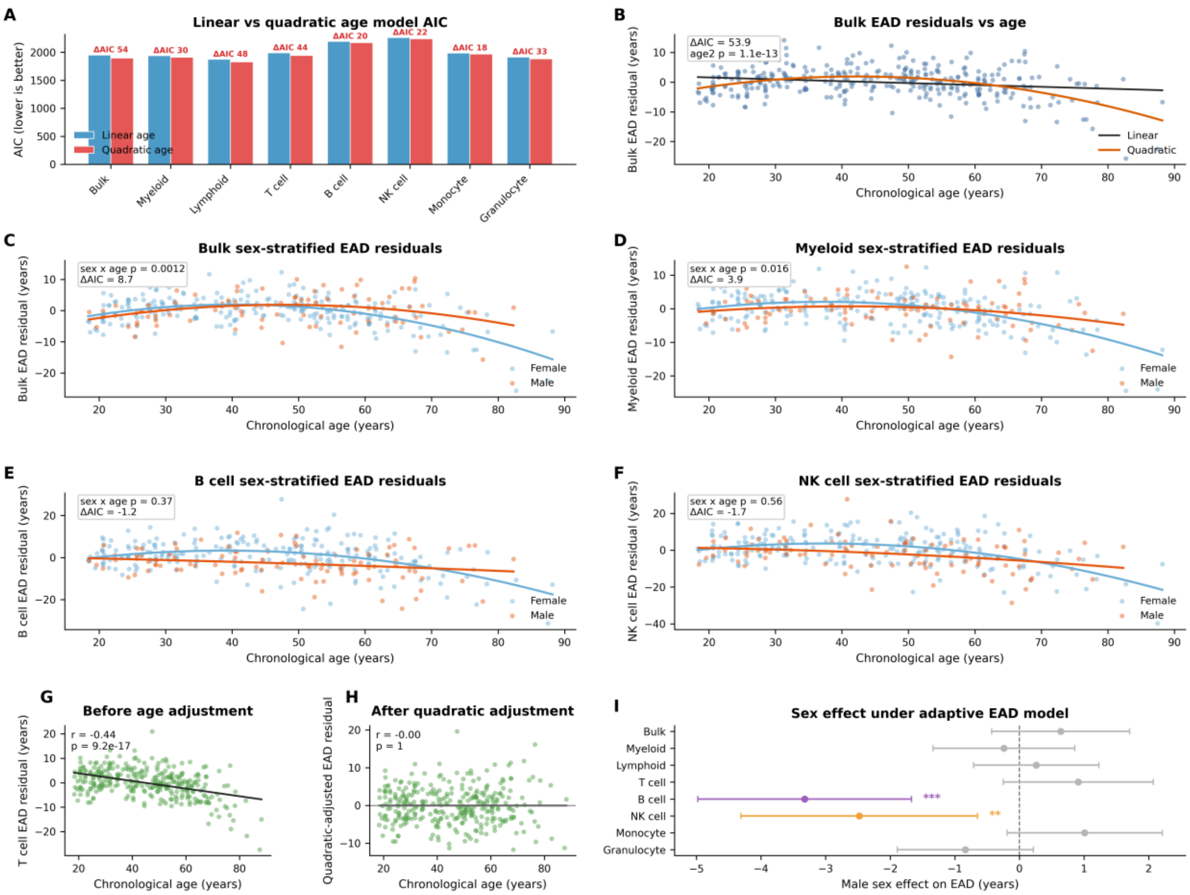


**Supplementary Figure S6.** Justification for quadratic age covariate and adaptive sex model in EAD association analyses. (a) AIC comparison between linear and quadratic age models across all eight clocks; the quadratic model is consistently preferred (DeltaAIC = 18–54). (b) Bulk clock EAD residuals plotted against chronological age with linear and quadratic fitted curves, illustrating the nonlinear age–residual relationship. (c–f) Sex-stratified EAD residuals as a function of age for representative clocks, showing age-dependent sex effects in myeloid-lineage clocks (panels c–d; bulk and myeloid) and age-constant effects in lymphoid-lineage clocks (panels e–f; B cell and NK cell). (g–h) T cell EAD residuals before and after quadratic age adjustment, confirming removal of residual age correlation. (i) Forest plot of sex effect on EAD across all eight clocks under the adaptive sex model.

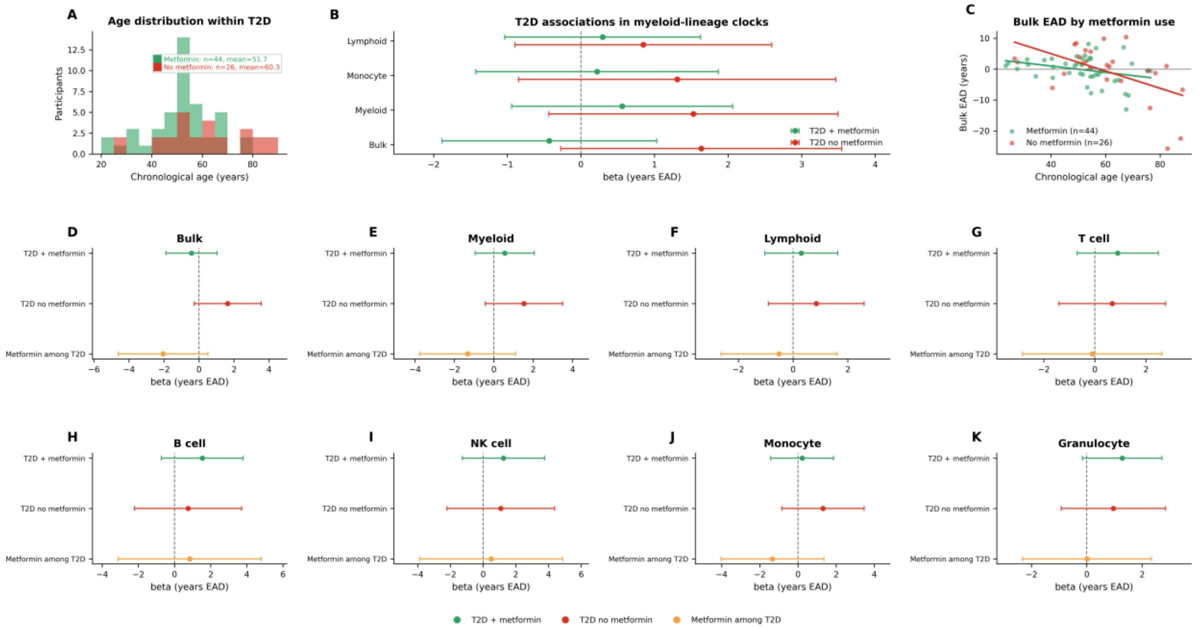


**Supplementary Figure S7.** Association of type 2 diabetes and metformin use with EAD across all eight cell-type clocks. Each panel shows three comparisons for one clock: T2D patients on metformin versus healthy controls (green), T2D patients not on metformin versus healthy controls (red), and metformin versus no metformin within T2D patients (orange). No comparison reached nominal significance ($p < 0.05$), likely reflecting limited power (n = 44 metformin, n = 26 untreated T2D, n = 252 healthy controls). The within-T2D comparison is additionally confounded by a mean age difference of 8.6 years between metformin-treated and untreated patients.

### Supplementary Tables

**Supplementary Table S1.** Full association results for ONT LongReadClock epigenetic age deviation across 31 phenotypes.

| Category | Phenotype | Clock | n | Cases | beta | 95% CI | p | q phenotype | q all |
|---|---|---|---|---|---|---|---|---|---|
| **Demographic** | | | | | | | | | |
| | Male sex | Bulk | 324 | 96 | +0.64 | [-0.43, +1.71] | 0.241 | 0.321 | 0.725 |
| | Male sex | Myeloid | 324 | 96 | -0.24 | [-1.34, +0.86] | 0.668 | 0.668 | 0.916 |
| | Male sex | Lymphoid | 324 | 96 | +0.26 | [-0.71, +1.23] | 0.598 | 0.668 | 0.901 |
| | Male sex | T cell | 324 | 96 | +0.91 | [-0.25, +2.07] | 0.124 | 0.198 | 0.631 |
| | Male sex | B cell | 324 | 96 | -3.32 | [-4.97, -1.68] | 9.38e-05 | 0.001 | 0.023 |
| | Male sex | NK cell | 324 | 96 | -2.48 | [-4.3, -0.65] | 0.008 | 0.033 | 0.271 |
| | Male sex | Monocyte | 324 | 96 | +1.01 | [-0.19, +2.21] | 0.099 | 0.198 | 0.611 |
| | Male sex | Granulocyte | 324 | 96 | -0.84 | [-1.89, +0.22] | 0.119 | 0.198 | 0.627 |
| **Lifestyle** | | | | | | | | | |
| | Ever heavy smoker | Bulk | 310 | 113 | +0.6 | [-0.47, +1.68] | 0.273 | 0.316 | 0.725 |
| | Ever heavy smoker | Myeloid | 310 | 113 | +0.75 | [-0.35, +1.85] | 0.181 | 0.316 | 0.639 |
| | Ever heavy smoker | Lymphoid | 310 | 113 | +0.54 | [-0.44, +1.52] | 0.277 | 0.316 | 0.725 |
| | Ever heavy smoker | T cell | 310 | 113 | +0.57 | [-0.6, +1.75] | 0.339 | 0.339 | 0.771 |
| | Ever heavy smoker | B cell | 310 | 113 | +1.53 | [-0.13, +3.18] | 0.071 | 0.316 | 0.536 |
| | Ever heavy smoker | NK cell | 310 | 113 | +1.04 | [-0.8, +2.88] | 0.268 | 0.316 | 0.725 |
| | Ever heavy smoker | Monocyte | 310 | 113 | +0.88 | [-0.32, +2.07] | 0.151 | 0.316 | 0.636 |
| | Ever heavy smoker | Granulocyte | 310 | 113 | +0.88 | [-0.18, +1.93] | 0.103 | 0.316 | 0.611 |
| | Current smoker | Bulk | 257 | 6 | +0.48 | [-0.83, +1.79] | 0.472 | 0.977 | 0.879 |
| | Current smoker | Myeloid | 257 | 6 | +0.07 | [-1.29, +1.44] | 0.915 | 0.977 | 0.98 |
| | Current smoker | Lymphoid | 257 | 6 | +0.42 | [-0.75, +1.59] | 0.482 | 0.977 | 0.881 |
| | Current smoker | T cell | 257 | 6 | +0.45 | [-0.94, +1.84] | 0.529 | 0.977 | 0.901 |
| | Current smoker | B cell | 257 | 6 | +0.17 | [-1.81, +2.14] | 0.87 | 0.977 | 0.976 |
| | Current smoker | NK cell | 257 | 6 | +0.03 | [-2.1, +2.17] | 0.977 | 0.977 | 0.995 |
| | Current smoker | Monocyte | 257 | 6 | +0.42 | [-1.07, +1.9] | 0.583 | 0.977 | 0.901 |
| | Current smoker | Granulocyte | 257 | 6 | +0.32 | [-0.99, +1.62] | 0.636 | 0.977 | 0.916 |
| | Current non-drinker | Bulk | 304 | 116 | -0.31 | [-1.35, +0.74] | 0.566 | 0.753 | 0.901 |
| | Current non-drinker | Myeloid | 304 | 116 | -0.5 | [-1.58, +0.58] | 0.365 | 0.753 | 0.794 |
| | Current non-drinker | Lymphoid | 304 | 116 | -0.39 | [-1.34, +0.56] | 0.423 | 0.753 | 0.826 |
| | Current non-drinker | T cell | 304 | 116 | -0.21 | [-1.35, +0.93] | 0.723 | 0.753 | 0.916 |
| | Current non-drinker | B cell | 304 | 116 | +0.45 | [-1.17, +2.07] | 0.586 | 0.753 | 0.901 |
| | Current non-drinker | NK cell | 304 | 116 | +0.29 | [-1.49, +2.07] | 0.753 | 0.753 | 0.916 |
| | Current non-drinker | Monocyte | 304 | 116 | -0.8 | [-1.98, +0.39] | 0.188 | 0.753 | 0.639 |
| | Current non-drinker | Granulocyte | 304 | 116 | -0.44 | [-1.48, +0.6] | 0.404 | 0.753 | 0.807 |
| **Physical health** | | | | | | | | | |
| | Difficulty walking/stairs | Bulk | 248 | 48 | +0.43 | [-1.08, +1.95] | 0.577 | 0.593 | 0.901 |

| Category | Phenotype | Clock | n | Cases | beta | 95% CI | p | q phenotype | q all |
|---|---|---|---|---|---|---|---|---|---|
| | Difficulty walking/stairs | Myeloid | 248 | 48 | +1.01 | [-0.56, +2.58] | 0.209 | 0.556 | 0.674 |
| | Difficulty walking/stairs | Lymphoid | 248 | 48 | +0.52 | [-0.88, +1.92] | 0.469 | 0.593 | 0.879 |
| | Difficulty walking/stairs | T cell | 248 | 48 | +0.45 | [-1.2, +2.11] | 0.593 | 0.593 | 0.901 |
| | Difficulty walking/stairs | B cell | 248 | 48 | +1.11 | [-1.2, +3.42] | 0.348 | 0.556 | 0.771 |
| | Difficulty walking/stairs | NK cell | 248 | 48 | +1.41 | [-1.16, +3.99] | 0.284 | 0.556 | 0.726 |
| | Difficulty walking/stairs | Monocyte | 248 | 48 | +1.6 | [-0.08, +3.28] | 0.063 | 0.502 | 0.536 |
| | Difficulty walking/stairs | Granulocyte | 248 | 48 | +1.01 | [-0.49, +2.51] | 0.187 | 0.556 | 0.639 |
| **Physiological** | | | | | | | | | |
| | Resting heart rate | Bulk | 321 | | +0.03 | [-0.02, +0.07] | 0.222 | 0.296 | 0.689 |
| | Resting heart rate | Myeloid | 321 | | +0.05 | [+0, +0.09] | 0.03 | 0.167 | 0.418 |
| | Resting heart rate | Lymphoid | 321 | | +0.02 | [-0.02, +0.06] | 0.381 | 0.436 | 0.807 |
| | Resting heart rate | T cell | 321 | | +0 | [-0.04, +0.05] | 0.922 | 0.922 | 0.98 |
| | Resting heart rate | B cell | 321 | | +0.04 | [-0.02, +0.11] | 0.213 | 0.296 | 0.676 |
| | Resting heart rate | NK cell | 321 | | +0.05 | [-0.02, +0.12] | 0.174 | 0.296 | 0.639 |
| | Resting heart rate | Monocyte | 321 | | +0.04 | [-0, +0.09] | 0.062 | 0.167 | 0.536 |
| | Resting heart rate | Granulocyte | 321 | | +0.04 | [-0, +0.08] | 0.063 | 0.167 | 0.536 |
| **Physical health** | | | | | | | | | |
| | Poor/fair self-rated health | Bulk | 316 | 108 | +0.22 | [-0.86, +1.29] | 0.695 | 0.932 | 0.916 |
| | Poor/fair self-rated health | Myeloid | 316 | 108 | +0.23 | [-0.88, +1.33] | 0.685 | 0.932 | 0.916 |
| | Poor/fair self-rated health | Lymphoid | 316 | 108 | +0.19 | [-0.79, +1.17] | 0.699 | 0.932 | 0.916 |
| | Poor/fair self-rated health | T cell | 316 | 108 | +0.01 | [-1.16, +1.17] | 0.991 | 0.991 | 0.995 |
| | Poor/fair self-rated health | B cell | 316 | 108 | +0.15 | [-1.48, +1.77] | 0.861 | 0.984 | 0.974 |
| | Poor/fair self-rated health | NK cell | 316 | 108 | -1.13 | [-2.93, +0.67] | 0.219 | 0.878 | 0.689 |
| | Poor/fair self-rated health | Monocyte | 316 | 108 | +0.32 | [-0.88, +1.53] | 0.6 | 0.932 | 0.901 |
| | Poor/fair self-rated health | Granulocyte | 316 | 108 | +0.75 | [-0.3, +1.81] | 0.164 | 0.878 | 0.636 |
| **Physiological** | | | | | | | | | |
| | BMI | Bulk | 321 | | -0.01 | [-0.08, +0.05] | 0.706 | 0.806 | 0.916 |
| | BMI | Myeloid | 321 | | +0.02 | [-0.05, +0.09] | 0.611 | 0.806 | 0.907 |
| | BMI | Lymphoid | 321 | | +0.02 | [-0.04, +0.08] | 0.488 | 0.806 | 0.881 |
| | BMI | T cell | 321 | | +0.03 | [-0.04, +0.11] | 0.386 | 0.806 | 0.807 |
| | BMI | B cell | 321 | | +0.12 | [+0.02, +0.23] | 0.025 | 0.196 | 0.38 |
| | BMI | NK cell | 321 | | +0.03 | [-0.08, +0.15] | 0.581 | 0.806 | 0.901 |
| | BMI | Monocyte | 321 | | -0.01 | [-0.08, +0.07] | 0.88 | 0.88 | 0.978 |
| | BMI | Granulocyte | 321 | | +0.06 | [-0.01, +0.13] | 0.082 | 0.329 | 0.551 |
| | Systolic blood pressure | Bulk | 321 | | +0.02 | [-0.02, +0.05] | 0.341 | 0.39 | 0.771 |
| | Systolic blood pressure | Myeloid | 321 | | +0.03 | [+0, +0.07] | 0.036 | 0.145 | 0.431 |
| | Systolic blood pressure | Lymphoid | 321 | | +0.02 | [-0.01, +0.05] | 0.188 | 0.3 | 0.639 |
| | Systolic blood pressure | T cell | 321 | | +0.03 | [-0.01, +0.06] | 0.146 | 0.292 | 0.636 |
| | Systolic blood pressure | B cell | 321 | | +0.03 | [-0.02, +0.08] | 0.279 | 0.372 | 0.725 |
| | Systolic blood pressure | NK cell | 321 | | +0.02 | [-0.03, +0.08] | 0.466 | 0.466 | 0.879 |
| | Systolic blood pressure | Monocyte | 321 | | +0.03 | [-0.01, +0.06] | 0.127 | 0.292 | 0.632 |

| Category | Phenotype | Clock | n | Cases | beta | 95% CI | p | q phenotype | q all |
|---|---|---|---|---|---|---|---|---|---|
| | Systolic blood pressure | Granulocyte | 321 | | +0.03 | [+0, +0.06] | 0.036 | 0.145 | 0.431 |
| **Medication** | | | | | | | | | |
| | Metformin use | Bulk | 324 | 46 | -0.48 | [-1.9, +0.94] | 0.51 | 0.779 | 0.897 |
| | Metformin use | Myeloid | 324 | 46 | +0.4 | [-1.06, +1.86] | 0.59 | 0.779 | 0.901 |
| | Metformin use | Lymphoid | 324 | 46 | +0.27 | [-1.02, +1.56] | 0.682 | 0.779 | 0.916 |
| | Metformin use | T cell | 324 | 46 | +0.74 | [-0.8, +2.29] | 0.346 | 0.697 | 0.771 |
| | Metformin use | B cell | 324 | 46 | +1.63 | [-0.56, +3.82] | 0.146 | 0.583 | 0.636 |
| | Metformin use | NK cell | 324 | 46 | +1.17 | [-1.27, +3.6] | 0.348 | 0.697 | 0.771 |
| | Metformin use | Monocyte | 324 | 46 | +0.09 | [-1.51, +1.69] | 0.912 | 0.912 | 0.98 |
| | Metformin use | Granulocyte | 324 | 46 | +1.17 | [-0.23, +2.56] | 0.101 | 0.583 | 0.611 |
| | Statin use | Bulk | 324 | 6 | -1.2 | [-2.53, +0.13] | 0.079 | 0.228 | 0.545 |
| | Statin use | Myeloid | 324 | 6 | -0.98 | [-2.35, +0.39] | 0.164 | 0.262 | 0.636 |
| | Statin use | Lymphoid | 324 | 6 | -0.88 | [-2.1, +0.33] | 0.155 | 0.262 | 0.636 |
| | Statin use | T cell | 324 | 6 | -1.35 | [-2.8, +0.1] | 0.068 | 0.228 | 0.536 |
| | Statin use | B cell | 324 | 6 | -0.26 | [-2.33, +1.82] | 0.807 | 0.807 | 0.948 |
| | Statin use | NK cell | 324 | 6 | -0.42 | [-2.72, +1.88] | 0.719 | 0.807 | 0.916 |
| | Statin use | Monocyte | 324 | 6 | -1.32 | [-2.82, +0.18] | 0.086 | 0.228 | 0.558 |
| | Statin use | Granulocyte | 324 | 6 | -0.48 | [-1.8, +0.84] | 0.476 | 0.635 | 0.881 |
| | Chronic steroid use | Bulk | 324 | 33 | +0.71 | [-0.9, +2.31] | 0.39 | 0.973 | 0.807 |
| | Chronic steroid use | Myeloid | 324 | 33 | -0.22 | [-1.87, +1.43] | 0.797 | 0.973 | 0.948 |
| | Chronic steroid use | Lymphoid | 324 | 33 | -0.08 | [-1.54, +1.39] | 0.919 | 0.973 | 0.98 |
| | Chronic steroid use | T cell | 324 | 33 | -0.5 | [-2.25, +1.24] | 0.572 | 0.973 | 0.901 |
| | Chronic steroid use | B cell | 324 | 33 | -0.13 | [-2.62, +2.36] | 0.917 | 0.973 | 0.98 |
| | Chronic steroid use | NK cell | 324 | 33 | -0.46 | [-3.21, +2.3] | 0.746 | 0.973 | 0.916 |
| | Chronic steroid use | Monocyte | 324 | 33 | +0.03 | [-1.78, +1.84] | 0.973 | 0.973 | 0.995 |
| | Chronic steroid use | Granulocyte | 324 | 33 | -0.46 | [-2.05, +1.12] | 0.567 | 0.973 | 0.901 |
| **Condition** | | | | | | | | | |
| | Hypertension | Bulk | 324 | 96 | -0.24 | [-1.38, +0.9] | 0.685 | 0.849 | 0.916 |
| | Hypertension | Myeloid | 324 | 96 | -0.05 | [-1.22, +1.13] | 0.94 | 0.94 | 0.98 |
| | Hypertension | Lymphoid | 324 | 96 | -0.53 | [-1.56, +0.5] | 0.313 | 0.849 | 0.753 |
| | Hypertension | T cell | 324 | 96 | -0.81 | [-2.05, +0.42] | 0.196 | 0.849 | 0.65 |
| | Hypertension | B cell | 324 | 96 | -0.68 | [-2.44, +1.08] | 0.449 | 0.849 | 0.863 |
| | Hypertension | NK cell | 324 | 96 | -0.57 | [-2.51, +1.38] | 0.569 | 0.849 | 0.901 |
| | Hypertension | Monocyte | 324 | 96 | -0.24 | [-1.52, +1.04] | 0.716 | 0.849 | 0.916 |
| | Hypertension | Granulocyte | 324 | 96 | +0.19 | [-0.94, +1.31] | 0.743 | 0.849 | 0.916 |
| | Obesity | Bulk | 324 | 18 | -0.01 | [-1, +0.99] | 0.988 | 0.995 | 0.995 |
| | Obesity | Myeloid | 324 | 18 | +0.2 | [-0.83, +1.22] | 0.707 | 0.943 | 0.916 |
| | Obesity | Lymphoid | 324 | 18 | +0.22 | [-0.69, +1.12] | 0.641 | 0.943 | 0.916 |
| | Obesity | T cell | 324 | 18 | +0.69 | [-0.39, +1.78] | 0.209 | 0.558 | 0.674 |
| | Obesity | B cell | 324 | 18 | +1.33 | [-0.2, +2.87] | 0.09 | 0.558 | 0.572 |

| Category | Phenotype | Clock | n | Cases | beta | 95% CI | p | q phenotype | q all |
|---|---|---|---|---|---|---|---|---|---|
| | Obesity | NK cell | 324 | 18 | -0.01 | [-1.72, +1.7] | 0.995 | 0.995 | 0.995 |
| | Obesity | Monocyte | 324 | 18 | +0.25 | [-0.87, +1.36] | 0.667 | 0.943 | 0.916 |
| | Obesity | Granulocyte | 324 | 18 | +0.73 | [-0.24, +1.71] | 0.142 | 0.558 | 0.636 |
| | Type 2 diabetes | Bulk | 324 | 7 | +0.29 | [-0.96, +1.53] | 0.653 | 0.653 | 0.916 |
| | Type 2 diabetes | Myeloid | 324 | 7 | +0.9 | [-0.38, +2.17] | 0.169 | 0.436 | 0.639 |
| | Type 2 diabetes | Lymphoid | 324 | 7 | +0.49 | [-0.64, +1.61] | 0.397 | 0.46 | 0.807 |
| | Type 2 diabetes | T cell | 324 | 7 | +0.82 | [-0.52, +2.17] | 0.232 | 0.436 | 0.709 |
| | Type 2 diabetes | B cell | 324 | 7 | +1.26 | [-0.65, +3.18] | 0.196 | 0.436 | 0.65 |
| | Type 2 diabetes | NK cell | 324 | 7 | +1.19 | [-0.93, +3.31] | 0.272 | 0.436 | 0.725 |
| | Type 2 diabetes | Monocyte | 324 | 7 | +0.6 | [-0.8, +1.99] | 0.403 | 0.46 | 0.807 |
| | Type 2 diabetes | Granulocyte | 324 | 7 | +1.17 | [-0.05, +2.39] | 0.061 | 0.436 | 0.536 |
| | Anxiety disorder | Bulk | 324 | 77 | +0.03 | [-1.11, +1.17] | 0.961 | 0.961 | 0.993 |
| | Anxiety disorder | Myeloid | 324 | 77 | -0.5 | [-1.67, +0.67] | 0.402 | 0.961 | 0.807 |
| | Anxiety disorder | Lymphoid | 324 | 77 | +0.04 | [-1, +1.08] | 0.94 | 0.961 | 0.98 |
| | Anxiety disorder | T cell | 324 | 77 | -0.09 | [-1.34, +1.15] | 0.884 | 0.961 | 0.978 |
| | Anxiety disorder | B cell | 324 | 77 | +0.07 | [-1.7, +1.84] | 0.941 | 0.961 | 0.98 |
| | Anxiety disorder | NK cell | 324 | 77 | +0.28 | [-1.68, +2.25] | 0.776 | 0.961 | 0.937 |
| | Anxiety disorder | Monocyte | 324 | 77 | -0.39 | [-1.67, +0.9] | 0.556 | 0.961 | 0.901 |
| | Anxiety disorder | Granulocyte | 324 | 77 | -0.24 | [-1.37, +0.88] | 0.674 | 0.961 | 0.916 |
| | Depression | Bulk | 324 | 6 | -0.44 | [-1.7, +0.81] | 0.489 | 0.95 | 0.881 |
| | Depression | Myeloid | 324 | 6 | -0.89 | [-2.18, +0.4] | 0.177 | 0.707 | 0.639 |
| | Depression | Lymphoid | 324 | 6 | -0.07 | [-1.21, +1.08] | 0.911 | 0.95 | 0.98 |
| | Depression | T cell | 324 | 6 | -0.24 | [-1.62, +1.13] | 0.727 | 0.95 | 0.916 |
| | Depression | B cell | 324 | 6 | -0.36 | [-2.31, +1.59] | 0.721 | 0.95 | 0.916 |
| | Depression | NK cell | 324 | 6 | -0.92 | [-3.09, +1.24] | 0.402 | 0.95 | 0.807 |
| | Depression | Monocyte | 324 | 6 | -1.03 | [-2.44, +0.38] | 0.155 | 0.707 | 0.636 |
| | Depression | Granulocyte | 324 | 6 | -0.04 | [-1.28, +1.2] | 0.95 | 0.95 | 0.986 |
| | Asthma | Bulk | 324 | 47 | +0.58 | [-0.79, +1.95] | 0.409 | 0.654 | 0.811 |
| | Asthma | Myeloid | 324 | 47 | -0.08 | [-1.5, +1.33] | 0.906 | 0.933 | 0.98 |
| | Asthma | Lymphoid | 324 | 47 | +0.55 | [-0.7, +1.8] | 0.388 | 0.654 | 0.807 |
| | Asthma | T cell | 324 | 47 | +0.82 | [-0.67, +2.32] | 0.281 | 0.654 | 0.725 |
| | Asthma | B cell | 324 | 47 | +1.7 | [-0.42, +3.83] | 0.117 | 0.467 | 0.627 |
| | Asthma | NK cell | 324 | 47 | +2.51 | [+0.17, +4.86] | 0.037 | 0.292 | 0.431 |
| | Asthma | Monocyte | 324 | 47 | +0.19 | [-1.35, +1.74] | 0.807 | 0.933 | 0.948 |
| | Asthma | Granulocyte | 324 | 47 | +0.06 | [-1.3, +1.41] | 0.933 | 0.933 | 0.98 |
| | NAFLD/fatty liver | Bulk | 324 | 43 | -1.31 | [-2.75, +0.14] | 0.077 | 0.441 | 0.545 |
| | NAFLD/fatty liver | Myeloid | 324 | 43 | -1.21 | [-2.7, +0.27] | 0.11 | 0.441 | 0.621 |
| | NAFLD/fatty liver | Lymphoid | 324 | 43 | -0.74 | [-2.06, +0.58] | 0.273 | 0.491 | 0.725 |
| | NAFLD/fatty liver | T cell | 324 | 43 | -0.27 | [-1.85, +1.32] | 0.74 | 0.846 | 0.916 |
| | NAFLD/fatty liver | B cell | 324 | 43 | +0.01 | [-2.24, +2.27] | 0.991 | 0.991 | 0.995 |

| Category | Phenotype | Clock | n | Cases | beta | 95% CI | p | q phenotype | q all |
|---|---|---|---|---|---|---|---|---|---|
| | NAFLD/fatty liver | NK cell | 324 | 43 | -0.43 | [-2.93, +2.06] | 0.734 | 0.846 | 0.916 |
| | NAFLD/fatty liver | Monocyte | 324 | 43 | -0.85 | [-2.48, +0.78] | 0.307 | 0.491 | 0.746 |
| | NAFLD/fatty liver | Granulocyte | 324 | 43 | -0.85 | [-2.28, +0.58] | 0.245 | 0.491 | 0.725 |
| | Chronic kidney disease | Bulk | 324 | 22 | -0.15 | [-2.15, +1.84] | 0.88 | 0.88 | 0.978 |
| | Chronic kidney disease | Myeloid | 324 | 22 | +0.54 | [-1.51, +2.6] | 0.604 | 0.88 | 0.902 |
| | Chronic kidney disease | Lymphoid | 324 | 22 | -0.57 | [-2.38, +1.24] | 0.539 | 0.88 | 0.901 |
| | Chronic kidney disease | T cell | 324 | 22 | -1.14 | [-3.3, +1.03] | 0.303 | 0.809 | 0.746 |
| | Chronic kidney disease | B cell | 324 | 22 | -2.24 | [-5.31, +0.84] | 0.155 | 0.809 | 0.636 |
| | Chronic kidney disease | NK cell | 324 | 22 | -1.95 | [-5.37, +1.46] | 0.263 | 0.809 | 0.725 |
| | Chronic kidney disease | Monocyte | 324 | 22 | -0.23 | [-2.47, +2.02] | 0.844 | 0.88 | 0.969 |
| | Chronic kidney disease | Granulocyte | 324 | 22 | +0.33 | [-1.64, +2.31] | 0.74 | 0.88 | 0.916 |
| | Coronary artery disease | Bulk | 324 | 19 | -0.46 | [-2.65, +1.72] | 0.679 | 0.911 | 0.916 |
| | Coronary artery disease | Myeloid | 324 | 19 | +0.31 | [-1.93, +2.56] | 0.787 | 0.911 | 0.942 |
| | Coronary artery disease | Lymphoid | 324 | 19 | -0.21 | [-2.19, +1.77] | 0.836 | 0.911 | 0.964 |
| | Coronary artery disease | T cell | 324 | 19 | -0.38 | [-2.76, +1.99] | 0.751 | 0.911 | 0.916 |
| | Coronary artery disease | B cell | 324 | 19 | +0.94 | [-2.43, +4.32] | 0.584 | 0.911 | 0.901 |
| | Coronary artery disease | NK cell | 324 | 19 | +0.62 | [-3.12, +4.36] | 0.745 | 0.911 | 0.916 |
| | Coronary artery disease | Monocyte | 324 | 19 | -0.14 | [-2.6, +2.32] | 0.911 | 0.911 | 0.98 |
| | Coronary artery disease | Granulocyte | 324 | 19 | +0.59 | [-1.57, +2.75] | 0.592 | 0.911 | 0.901 |
| | Myocardial infarction | Bulk | 324 | 15 | -1.75 | [-4.22, +0.71] | 0.164 | 0.368 | 0.636 |
| | Myocardial infarction | Myeloid | 324 | 15 | -0.77 | [-3.31, +1.77] | 0.553 | 0.593 | 0.901 |
| | Myocardial infarction | Lymphoid | 324 | 15 | -1.49 | [-3.69, +0.7] | 0.184 | 0.368 | 0.639 |
| | Myocardial infarction | T cell | 324 | 15 | -1.98 | [-4.6, +0.65] | 0.141 | 0.368 | 0.636 |
| | Myocardial infarction | B cell | 324 | 15 | -3.66 | [-7.38, +0.07] | 0.055 | 0.368 | 0.536 |
| | Myocardial infarction | NK cell | 324 | 15 | -1.13 | [-5.28, +3.02] | 0.593 | 0.593 | 0.901 |
| | Myocardial infarction | Monocyte | 324 | 15 | -0.97 | [-3.75, +1.81] | 0.496 | 0.593 | 0.883 |
| | Myocardial infarction | Granulocyte | 324 | 15 | -0.86 | [-3.3, +1.58] | 0.49 | 0.593 | 0.881 |
| **Physiological** | | | | | | | | | |
| | Waist circumference | Bulk | 260 | | +0.01 | [-0.03, +0.04] | 0.71 | 0.778 | 0.916 |
| | Waist circumference | Myeloid | 260 | | +0.02 | [-0.02, +0.05] | 0.274 | 0.483 | 0.725 |
| | Waist circumference | Lymphoid | 260 | | +0.02 | [-0.01, +0.05] | 0.258 | 0.483 | 0.725 |
| | Waist circumference | T cell | 260 | | +0.02 | [-0.02, +0.06] | 0.328 | 0.483 | 0.768 |
| | Waist circumference | B cell | 260 | | +0.05 | [-0, +0.11] | 0.071 | 0.285 | 0.536 |
| | Waist circumference | NK cell | 260 | | +0.03 | [-0.03, +0.09] | 0.362 | 0.483 | 0.794 |
| | Waist circumference | Monocyte | 260 | | +0.01 | [-0.03, +0.04] | 0.778 | 0.778 | 0.937 |
| | Waist circumference | Granulocyte | 260 | | +0.03 | [-2.93e-05, +0.07] | 0.051 | 0.285 | 0.536 |
| **Laboratory** | | | | | | | | | |
| | HDL cholesterol | Bulk | 111 | | +0.01 | [-0.04, +0.05] | 0.814 | 0.985 | 0.948 |
| | HDL cholesterol | Myeloid | 111 | | -0.01 | [-0.06, +0.03] | 0.577 | 0.985 | 0.901 |
| | HDL cholesterol | Lymphoid | 111 | | +0 | [-0.04, +0.04] | 0.934 | 0.985 | 0.98 |

| Category | Phenotype | Clock | n | Cases | beta | 95% CI | p | q phenotype | q all |
|---|---|---|---|---|---|---|---|---|---|
| | HDL cholesterol | T cell | 111 | | -0 | [-0.05, +0.05] | 0.985 | 0.985 | 0.995 |
| | HDL cholesterol | B cell | 111 | | -0.06 | [-0.13, +0] | 0.057 | 0.229 | 0.536 |
| | HDL cholesterol | NK cell | 111 | | -0.09 | [-0.17, -0.02] | 0.02 | 0.161 | 0.332 |
| | HDL cholesterol | Monocyte | 111 | | -0 | [-0.05, +0.05] | 0.979 | 0.985 | 0.995 |
| | HDL cholesterol | Granulocyte | 111 | | -0.01 | [-0.06, +0.03] | 0.578 | 0.985 | 0.901 |
| | LDL cholesterol | Bulk | 107 | | -0.04 | [-0.07, -0.01] | 0.008 | 0.021 | 0.271 |
| | LDL cholesterol | Myeloid | 107 | | -0.03 | [-0.06, -0.01] | 0.017 | 0.033 | 0.307 |
| | LDL cholesterol | Lymphoid | 107 | | -0.04 | [-0.06, -0.01] | 0.002 | 0.019 | 0.227 |
| | LDL cholesterol | T cell | 107 | | -0.02 | [-0.05, +0.01] | 0.125 | 0.166 | 0.631 |
| | LDL cholesterol | B cell | 107 | | -0.02 | [-0.06, +0.02] | 0.317 | 0.317 | 0.755 |
| | LDL cholesterol | NK cell | 107 | | -0.03 | [-0.08, +0.02] | 0.261 | 0.298 | 0.725 |
| | LDL cholesterol | Monocyte | 107 | | -0.03 | [-0.06, +0] | 0.068 | 0.109 | 0.536 |
| | LDL cholesterol | Granulocyte | 107 | | -0.04 | [-0.06, -0.01] | 0.007 | 0.021 | 0.271 |
| | Triglycerides | Bulk | 106 | | -0.01 | [-0.02, +0.01] | 0.442 | 0.536 | 0.856 |
| | Triglycerides | Myeloid | 106 | | +0 | [-0.01, +0.02] | 0.469 | 0.536 | 0.879 |
| | Triglycerides | Lymphoid | 106 | | -0 | [-0.01, +0.01] | 0.719 | 0.719 | 0.916 |
| | Triglycerides | T cell | 106 | | -0.01 | [-0.02, +0.01] | 0.417 | 0.536 | 0.821 |
| | Triglycerides | B cell | 106 | | -0.01 | [-0.03, +0.01] | 0.298 | 0.536 | 0.745 |
| | Triglycerides | NK cell | 106 | | -0.03 | [-0.05, -0.01] | 0.017 | 0.139 | 0.307 |
| | Triglycerides | Monocyte | 106 | | +0.01 | [-0.01, +0.02] | 0.307 | 0.536 | 0.746 |
| | Triglycerides | Granulocyte | 106 | | +0.01 | [-0, +0.02] | 0.148 | 0.536 | 0.636 |
| | Total cholesterol | Bulk | 111 | | -0.03 | [-0.05, -0.01] | 0.016 | 0.042 | 0.307 |
| | Total cholesterol | Myeloid | 111 | | -0.02 | [-0.04, +0] | 0.113 | 0.151 | 0.623 |
| | Total cholesterol | Lymphoid | 111 | | -0.02 | [-0.04, -0.01] | 0.013 | 0.042 | 0.307 |
| | Total cholesterol | T cell | 111 | | -0.02 | [-0.05, +0] | 0.078 | 0.151 | 0.545 |
| | Total cholesterol | B cell | 111 | | -0.03 | [-0.06, +0.01] | 0.108 | 0.151 | 0.621 |
| | Total cholesterol | NK cell | 111 | | -0.05 | [-0.08, -0.01] | 0.015 | 0.042 | 0.307 |
| | Total cholesterol | Monocyte | 111 | | -0.01 | [-0.04, +0.01] | 0.247 | 0.247 | 0.725 |
| | Total cholesterol | Granulocyte | 111 | | -0.02 | [-0.04, +0.01] | 0.157 | 0.18 | 0.636 |
| | HbA1c | Bulk | 122 | | +0.06 | [-0.46, +0.57] | 0.83 | 0.83 | 0.962 |
| | HbA1c | Myeloid | 122 | | +0.28 | [-0.2, +0.77] | 0.257 | 0.389 | 0.725 |
| | HbA1c | Lymphoid | 122 | | +0.31 | [-0.12, +0.73] | 0.156 | 0.313 | 0.636 |
| | HbA1c | T cell | 122 | | +0.58 | [+0.07, +1.1] | 0.028 | 0.113 | 0.413 |
| | HbA1c | B cell | 122 | | +0.93 | [+0.25, +1.61] | 0.009 | 0.07 | 0.271 |
| | HbA1c | NK cell | 122 | | +0.27 | [-0.57, +1.1] | 0.533 | 0.609 | 0.901 |
| | HbA1c | Monocyte | 122 | | +0.29 | [-0.25, +0.84] | 0.291 | 0.389 | 0.738 |
| | HbA1c | Granulocyte | 122 | | +0.47 | [+0.01, +0.94] | 0.049 | 0.13 | 0.536 |
| | eGFR | Bulk | 78 | | -0.04 | [-0.08, -0.01] | 0.015 | 0.041 | 0.307 |
| | eGFR | Myeloid | 78 | | -0.02 | [-0.06, +0.01] | 0.179 | 0.359 | 0.639 |
| | eGFR | Lymphoid | 78 | | -0.04 | [-0.07, -0.01] | 0.004 | 0.015 | 0.227 |

| Category | Phenotype | Clock | n | Cases | beta | 95% CI | p | q phenotype | q all |
|---|---|---|---|---|---|---|---|---|---|
| | eGFR | T cell | 78 | | -0.05 | [-0.08, -0.02] | 0.003 | 0.015 | 0.227 |
| | eGFR | B cell | 78 | | -0.01 | [-0.06, +0.04] | 0.619 | 0.619 | 0.914 |
| | eGFR | NK cell | 78 | | -0.02 | [-0.07, +0.04] | 0.531 | 0.619 | 0.901 |
| | eGFR | Monocyte | 78 | | -0.01 | [-0.05, +0.03] | 0.556 | 0.619 | 0.901 |
| | eGFR | Granulocyte | 78 | | -0.02 | [-0.05, +0.02] | 0.32 | 0.513 | 0.757 |
| **Clinical burden** | | | | | | | | | |
| | Multimorbidity | Bulk | 324 | | +0 | [-0.01, +0.01] | 0.813 | 0.864 | 0.948 |
| | Multimorbidity | Myeloid | 324 | | -0 | [-0.01, +0.01] | 0.864 | 0.864 | 0.974 |
| | Multimorbidity | Lymphoid | 324 | | -0 | [-0.01, +0.01] | 0.702 | 0.864 | 0.916 |
| | Multimorbidity | T cell | 324 | | -0 | [-0.01, +0.01] | 0.685 | 0.864 | 0.916 |
| | Multimorbidity | B cell | 324 | | +0 | [-0.01, +0.02] | 0.86 | 0.864 | 0.974 |
| | Multimorbidity | NK cell | 324 | | +0 | [-0.01, +0.02] | 0.626 | 0.864 | 0.916 |
| | Multimorbidity | Monocyte | 324 | | -0.01 | [-0.02, +0.01] | 0.347 | 0.864 | 0.771 |
| | Multimorbidity | Granulocyte | 324 | | +0 | [-0.01, +0.01] | 0.811 | 0.864 | 0.948 |
| **Medication** | | | | | | | | | |
| | Polypharmacy | Bulk | 324 | | +0.01 | [-0.01, +0.02] | 0.401 | 0.903 | 0.807 |
| | Polypharmacy | Myeloid | 324 | | +0 | [-0.01, +0.01] | 0.852 | 0.903 | 0.974 |
| | Polypharmacy | Lymphoid | 324 | | +0 | [-0.01, +0.01] | 0.719 | 0.903 | 0.916 |
| | Polypharmacy | T cell | 324 | | +0 | [-0.01, +0.01] | 0.903 | 0.903 | 0.98 |
| | Polypharmacy | B cell | 324 | | -0 | [-0.02, +0.02] | 0.705 | 0.903 | 0.916 |
| | Polypharmacy | NK cell | 324 | | +0 | [-0.02, +0.02] | 0.732 | 0.903 | 0.916 |
| | Polypharmacy | Monocyte | 324 | | -0 | [-0.02, +0.01] | 0.686 | 0.903 | 0.916 |
| | Polypharmacy | Granulocyte | 324 | | +0 | [-0.01, +0.02] | 0.498 | 0.903 | 0.883 |

n is the complete-case sample size for each model. Cases are shown only for binary phenotypes. Beta is in years of EAD. q phenotype is Benjamini–Hochberg adjusted across the eight clocks within each phenotype; q all is adjusted across the full 31-phenotype screen. Obesity is defined as obesity diagnosis or BMI >= 30 kg/m2; HbA1c combines the standard total-blood and HPLC LOINC concepts.

**Supplementary Table S2.** Phenotype coverage for the ONT EAD association screen.

| Category | Phenotype | Present | Clock tests | Min n | Nominal hits |
|---|---|---|---|---|---|
| **Demographic** | | | | | |
| | Male sex | Yes | 8 | 324 | 2 |
| **Lifestyle** | | | | | |
| | Ever heavy smoker | Yes | 8 | 310 | 0 |
| | Current smoker | Yes | 8 | 257 | 0 |
| | Current non-drinker | Yes | 8 | 304 | 0 |
| **Physical health** | | | | | |
| | Difficulty walking/stairs | Yes | 8 | 248 | 0 |
| **Physiological** | | | | | |
| | Resting heart rate | Yes | 8 | 321 | 1 |
| **Physical health** | | | | | |
| | Poor/fair self-rated health | Yes | 8 | 316 | 0 |
| **Physiological** | | | | | |
| | BMI | Yes | 8 | 321 | 1 |
| | Systolic blood pressure | Yes | 8 | 321 | 2 |
| **Medication** | | | | | |
| | Metformin use | Yes | 8 | 324 | 0 |
| | Statin use | Yes | 8 | 324 | 0 |
| | Chronic steroid use | Yes | 8 | 324 | 0 |
| **Condition** | | | | | |
| | Hypertension | Yes | 8 | 324 | 0 |
| | Obesity | Yes | 8 | 324 | 0 |
| | Type 2 diabetes | Yes | 8 | 324 | 0 |
| | Anxiety disorder | Yes | 8 | 324 | 0 |
| | Depression | Yes | 8 | 324 | 0 |
| | Asthma | Yes | 8 | 324 | 1 |
| | NAFLD/fatty liver | Yes | 8 | 324 | 0 |
| | Chronic kidney disease | Yes | 8 | 324 | 0 |
| | Coronary artery disease | Yes | 8 | 324 | 0 |
| | Myocardial infarction | Yes | 8 | 324 | 0 |
| **Physiological** | | | | | |
| | Waist circumference | Yes | 8 | 260 | 0 |
| **Laboratory** | | | | | |
| | HDL cholesterol | Yes | 8 | 111 | 1 |
| | LDL cholesterol | Yes | 8 | 107 | 4 |
| | Triglycerides | Yes | 8 | 106 | 1 |
| | Total cholesterol | Yes | 8 | 111 | 3 |
| | HbA1c | Yes | 8 | 122 | 3 |
| | eGFR | Yes | 8 | 78 | 3 |
| **Clinical burden** | | | | | |
| | Multimorbidity | Yes | 8 | 324 | 0 |
| **Medication** | | | | | |
| | Polypharmacy | Yes | 8 | 324 | 0 |

Min n is the minimum complete-case sample size across the eight EAD clock models for the phenotype. Nominal hits are clock-specific associations with $p < 0.05$ before multiple-testing correction.
